# Sequential and coordinated control of human plasma cell differentiation by IRF4 and BLIMP1 associated with a discriminating ISRE/EICE motif lexicon

**DOI:** 10.64898/2026.06.15.732353

**Authors:** Louis C. Lau, Sneha Lal, Jingyu Fan, Nicholas A. Pease, Harinder Singh

## Abstract

The transcription factors IRF4 and BLIMP1 (*PRDM1*) promote plasma cell (PC) differentiation by repressing B cell identity genes while inducing the unfolded protein response, ER/Golgi biogenesis, and immunoglobulin secretion. How IRF4 and BLIMP1 partition and coordinate their genomic activities during plasma cell differentiation remains unresolved. Using naïve human B cells and a stepwise *in vitro* differentiation system, we performed CRISPR/Cas9 perturbations of *IRF4* or *PRDM1* in activated B-lineage cells that contained demonstrable plasma cell precursors, followed by single-cell RNA sequencing. Despite their mutually reinforced expression and shared recognition of related IRF-family motifs (ISREs and EICEs), loss of IRF4, but not of BLIMP1, yielded a stunted intermediate with incomplete silencing of B cell identity genes and defective induction of the secretory program. Multiome profiling, base-pair-resolution motif modeling, CUT&RUN, and DNA-binding assays identified non-conserved nucleotides within ISRE/EICE motifs that discriminate IRF4 from BLIMP1 binding. These findings define an ISRE/EICE motif lexicon that predicts differential IRF4 and BLIMP1 occupancy and are consistent with a temporally ordered model, in which IRF4 first acts independently and then in concert with BLIMP1 to promote PC differentiation.

## Introduction

Long-lived plasma cells (PCs) underpin durable protective humoral immune responses to pathogens and vaccines. Plasma cell differentiation involves dramatic chromatin and transcriptional remodeling that results in repression of an extensive set of B-lineage genes (Alaterre et al., 2024; Guo et al., 2018; Lio et al., 2016; Scharer et al., 2018; Shaffer et al., 2002; Shaknovich et al., 2011). The repression program is coupled with induction of a suite of genes regulating the unfolded protein response (UPR), endoplasmic reticulum (ER), and Golgi biogenesis to support high rates of antibody production and secretion (Nutt et al., 2015; Tellier et al., 2016; Wiggins & Scharer, 2021). The durability of this differentiated state is in turn sustained by metabolic remodeling and reprogramming of cell survival genes, resulting in a long-lived secretory state (Low et al., 2019). Although mature PCs have been extensively characterized, the stage-resolved regulatory logic that initiates and stabilizes terminal PC differentiation from a precursor (prePC) state remains to be elucidated.

Genetic analyses in mice have established that the transcription factors (TFs) IRF4 and BLIMP1 (*PRDM1*) are central determinants of the plasma cell fate. In activated murine B cells, graded IRF4 accumulation acts as a fate switch: lower IRF4 promotes germinal-center (GC) programming and recycling, whereas higher IRF4 promotes PC commitment and GC exit by engaging a transcriptional program that includes ZBTB20, BLIMP1, and downstream regulators such as XBP1 and ATF6B (Chevrier et al., 2014; Gloury et al., 2016; Ochiai et al., 2013; Sciammas et al., 2006; Shaffer et al., 2004; Xu et al., 2015). BLIMP1 is required to consolidate this transition by repressing B-lineage and proliferative gene programs (e.g., components of the IRF8, PAX5, EBF1, PU.1/SpiB, BCL6-driven B-cell identity circuit) while promoting terminal differentiation modules linked to immunoglobulin (Ig) secretion, the UPR, and ER/Golgi biogenesis (Conter et al., 2025; Deng et al., 2010; Minnich et al., 2016; Shapiro-Shelef et al., 2005; Tellier et al., 2016). Although BLIMP1 is often emphasized as the key effector of repression during B-to-PC differentiation, both IRF4 and BLIMP1 have been implicated in repression of B-cell identity during PC differentiation (Chevrier et al., 2014; Gloury et al., 2016; Shaffer et al., 2002; Wohner et al., 2016). Conditional perturbation of *Irf4* has also revealed its importance for maintenance of long-lived bone-marrow PCs (BMPCs), whereas conditional loss of *Prdm1* retains BMPCs but abolishes Ig secretory capacity (Low et al., 2019; Minnich et al., 2016; Shapiro-Shelef et al., 2005; Tellier et al., 2016). These observations suggest a temporally ordered interplay between IRF4 and BLIMP1, yet they do not resolve which components of the PC program are initiated by IRF4 before BLIMP1 accumulation, that are then consolidated by BLIMP1, and which genomic regions each TF targets within human prePCs.

Mechanistically, IRF4 and BLIMP1 converge on related *cis*-regulatory logic centered on their recognition of related DNA binding motifs (Kuo & Calame, 2004; Minnich et al., 2016). IRF4 is a structurally divergent, immune-restricted member of the interferon <u>r</u>egulatory factor (IRF) family that can engage DNA composite elements (CEs) through its interactions with distinct partners (Escalante et al., 2002; Honda & Taniguchi, 2006; Sundararaj et al., 2021). Classically these involve its interactions with ETS-family factors PU.1/SpiB at <u>E</u>TS–IRF <u>c</u>omposite <u>e</u>lements (EICEs) or with AP-1 family complexes at <u>A</u>P-1-IRF <u>c</u>omposite <u>e</u>lements (AICEs) (Brass et al., 1996; Escalante et al., 2002; Glasmacher et al., 2012; Ochiai et al., 2013; Sundararaj et al., 2021). Notably, the structurally related and immune-restricted IRF family TF, IRF8, displays similar DNA recognition specificity to IRF4 despite functioning to counteract IRF4 in promoting plasmablast (PB) and PC differentiation in activated B cells (Brass et al., 1996; Shukla & Lu, 2014; Xu et al., 2015). Importantly, IRF4 and IRF8 can also independently bind to canonical IRF-binding sites termed interferon-<u>s</u>timulated response <u>e</u>lements (ISREs) and antagonize the action of other IRF family members (Bovolenta et al., 1994; Honda & Taniguchi, 2006).

The EICEs appear to represent a subset of ISREs, as prototypical IRFs like IRF1 can bind EICEs efficiently as shown with the λB site in an Igλ enhancer (Brass et al., 1996). EICE-specific genomic regulation likely arose later than the canonical ISREs present in early metazoans, as IRF4/IRF8 and the ETS paralogs PU.1/SpiB arose from gene duplications with specialization in vertebrates after prototypical IRFs developed (Anderson et al., 2001; Nehyba et al., 2009; Shukla & Lu, 2014). BLIMP1 is a PR/SET-domain zinc-finger transcription factor that also has a later vertebrate evolutionary origin and binds subsets of ISREs and EICEs thereby antagonizing the action of prototypical IRFs such as IRF1, or immune-restricted IRFs IRF4/8 and the ETS family factors PU.1/SpiB, respectively (Minnich et al., 2016; Perdiguero et al., 2020; Shaffer et al., 2002). Importantly, subtle nucleotide differences within ISRE/EICE-like motifs have been suggested to bias binding of these TFs to ISREs or EICEs and regulatory outcomes, most notably BLIMP1’s specificity for sequences containing GT-GAAA-G (Doody et al., 2010; Minnich et al., 2016). However, genome-wide analyses of IRF4 and BLIMP1 binding at regulatory regions containing ISREs and/or EICEs, and the motif lexicon that underlies their independent and concerted functions in prePCs, remain to be explored. We sought to define this ISRE/EICE motif lexicon as the set of nucleotide-level variants within related IRF-family motifs that may differentially recruit IRF4 or BLIMP1 and thereby encode distinct regulatory outputs.

Here, we address these biological and molecular propositions by performing stage-specific perturbations of *IRF4* or *PRDM1* in human prePCs generated from primary naïve B cells using a stepwise *in vitro* differentiation system. We couple stage-specific Cas9 editing with single-cell RNA sequencing to resolve fate trajectories and stunted states. Furthermore, we integrate multiome profiling, base pair-resolution motif modeling, and CUT&RUN assays to connect the motif lexicon to factor occupancy and their regulatory outputs. Finally, by using electrophoretic mobility shift assays (EMSAs) to systematically test nucleotide determinants of IRF4 and BLIMP1 binding to ISREs and EICEs, we uncover the lexicon that enables both distinct and coordinated regulatory activity. The overall approach reveals how IRF4 and BLIMP1 differentially shape the prePC transcriptional program and identifies motif features that predict overlapping versus distinct genomic engagement. These observations are compatible with a temporally ordered interplay between IRF4 and BLIMP1 in human PC differentiation.

## Results

### A stepwise human B cell *in vitro* differentiation system generates plasma cell precursors that mature into PCs

To enable stage-specific perturbation of IRF4 and BLIMP1 during human PC differentiation, we used a stepwise *in vitro* culture system in which naïve B cells undergo activation and expansion, transition through a PB phase and undergo PC maturation (Figure 1A) (Cocco et al., 2012; Robinson et al., 2019; Su et al., 2016). We sampled cultures temporally (D0-21) and analyzed emergent cell populations by flow cytometry using established markers of PBs and PCs (Table S1) (Sanz et al., 2019; Staupe et al., 2022). CD27 and CD38 expression were up-regulated between D4 and D7 within a subset of the activated B cell population (actBs, CD20^+^CD38^+^CD27^-^) marking the emergence of PBs (CD20^-^CD38^+^CD27^+^CD138^-^) (Figure S1A). The D7 cultures were primarily composed of actBs and PBs (Figure 1B, C and S1B, C). After D7, PCs (CD20^-^ CD38^+^CD27^+^ICAM2^+^CD138^+^) began to emerge while PBs decreased, consistent with progressive maturation (Figure 1B, C and S1B, C). Notably, ICAM-2 upregulation preceded detectable CD138, providing an early surface marker for cells entering the PC trajectory in this system (Figure 1B and S1B), although ICAM-2 alone did not identify PCs.

**Figure 1.**
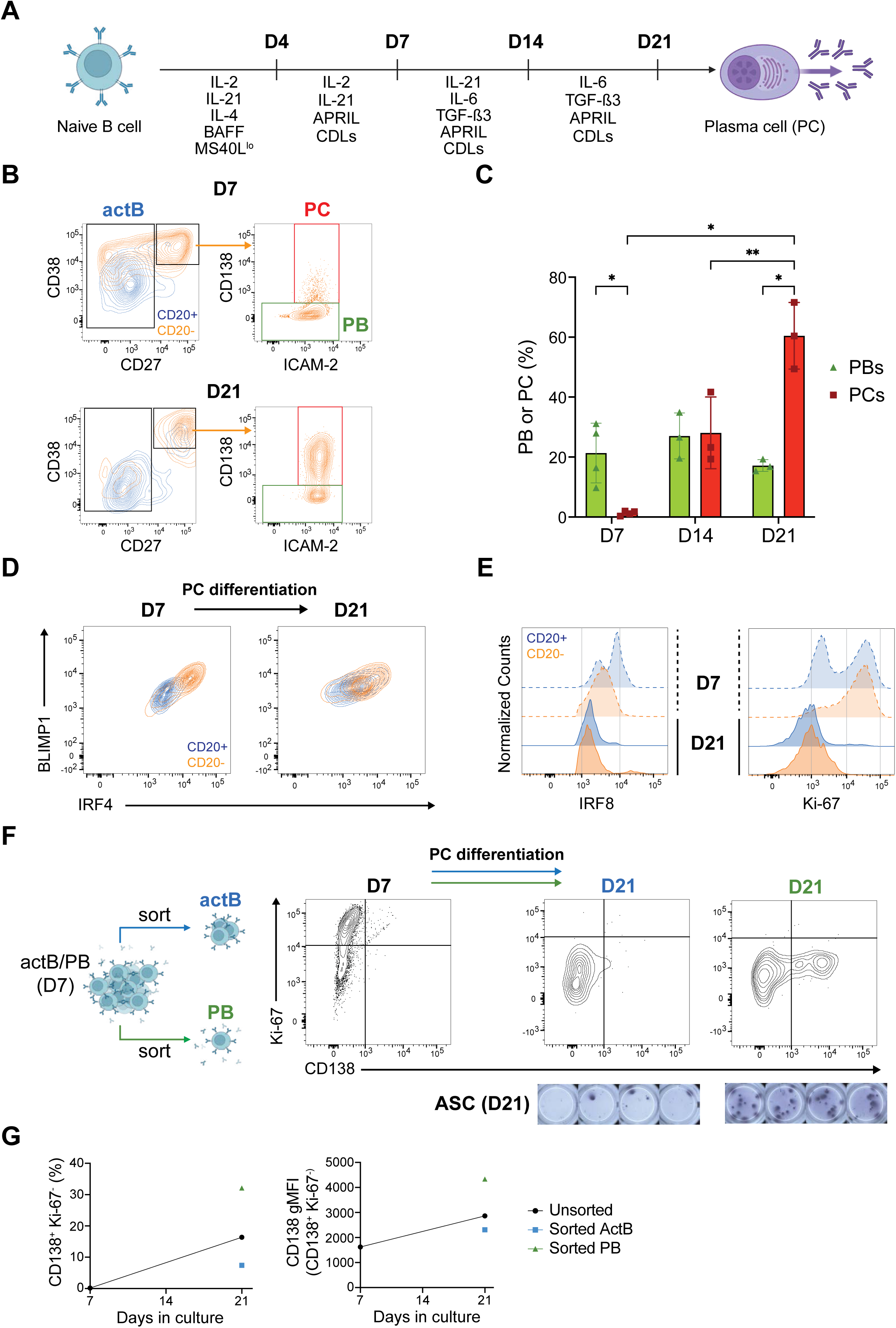
A stepwise human B cell *in vitro* differentiation system generates plasma cell precursors that mature into PCs. (A) Overview of the *in vitro* PC differentiation culture system. Naive human B cells isolated from healthy donor PBMCs were cultured through sequential phases of activation/expansion (D0-D4), PB/prePC generation (D4-D7), PC differentiation (D7-D14), and PC maturation/maintenance (D14+). (B, C) PB and PC differentiation were monitored by surface immunophenotyping. (B) Representative flow cytometry plots show gating strategies used to quantify actBs (CD20^+^CD38^+^CD27^-^), PBs (CD20^-^CD38^+^CD27^+^ICAM2^+/-^CD138^-^), and PCs (CD20^-^CD38^+^CD27^+^ICAM2^+^CD138^+^) at D7 and D21. (C) Bar graphs show PB and PC frequencies among live cells at the indicated timepoints (Donor 1). Dots indicate independent experiments, and error bars indicate mean +/- SD (n=3-4; *p<0.05, **p<0.01; two-way ANOVA with Tukey’s post-hoc test). (D, E) Intracellular flow cytometry was used to measure key transcription factors (BLIMP1, IRF4, IRF8) and proliferative status (Ki-67) along the PB/PC trajectory. To relate intracellular measurements to the surface-defined states in Figure 1B, cells were gated as CD20^+^ (blue) or CD20^-^ (orange) and displayed for the indicated proteins and timepoints. (D) Biplots showing BLIMP1 and IRF4 protein levels at D7 (left) and D21 (right). (E) Mode-normalized histograms of IRF8 and Ki-67 at D7 (dashed) and D21 (solid). (F) (Left) Schematic illustrates experimental design used to determine which D7 compartment contains functionally competent prePCs. (Right) The actB and PB populations were sorted at D7, placed back into identical D7-D21 culture conditions, and analyzed by flow cytometry and IgG ELISPOT at D21 (10% contour intervals; representative of 2 donors; ELISPOT: n=4 technical replicates; actB-generated input=375 cells; PB-generated input=236 cells). (G) Line graphs quantifying ICS biplots in (F), showing increased non-proliferative (Ki-67^-^) CD138^+^ cell frequency and CD138 gMFI in those cells at D21 in unsorted cultures. Additionally, re-cultured PBs have higher Ki-67^-^ CD138^+^ frequency and sub-gated CD138 expression compared to re-cultured actBs.

Next, we sought to map the expression of key transcription factors onto the PB/PC trajectory by linking intracellular measurements of IRF4, BLIMP1 and IRF8 with that of CD20, which demarcates a transition to the PB/PC trajectory and displays a strong correspondence between cell surface and intracellular staining protocols (Figure S1D). IRF4 and BLIMP1 proteins were induced during B cell activation and accumulated in two regulatory states as cells transitioned from actB cells (CD20^+^) to the PB/PC trajectory (CD20^-^). These were manifested as two regulatory regimes: 1) an IRF4^lo^BLIMP1^lo^ state and 2) an IRF4^hi^BLIMP1^hi^ state (Figure 1D and S1E). By D21, nearly all cells had exited the cell cycle (Ki-67^-^), fully downregulated IRF8, and maintained high expression levels of IRF4 and BLIMP1 consistent with terminal PC differentiation (Figure 1E and S1F). Thus, the cells at late timepoints in these cultures satisfied a comprehensive set of phenotypic and regulatory state criteria for mature PCs, including loss of CD20 and IRF8 and high expression of CD27, CD38, ICAM-2, CD138, IRF4 and BLIMP1 as well as cell-cycle withdrawal.

We next sought to define the population of cells containing PC precursors (prePCs) based on their functional competency in generating PCs. We considered two possibilities: in one scenario prePCs might derive from actB cells which switch from initially giving rise to PBs to differentiating into PCs. Conversely, prePCs could be contained within the canonical PB population i.e., CD20^-^CD38^+^CD27^+^CD138^-^ cells. To distinguish between these possibilities and directly compare their PC-generating competencies, we sorted D7 CD20^+^CD27^-^ actB and CD20^-^CD38^+^CD27^+^ PBs for re-culture under identical differentiation conditions between D7-D21, before measuring total or surface CD138 expression at D21 in the context of the entire staining panels (Figure 1F and S1G). In contrast to the sorted actB population which generated very few PCs at D21 (7±1 PC/10,000 actB), the sorted PB population showed an enhanced capacity to generate PCs at D21 (360±140 PC/10,000 PB; 50-fold increase; Table S1). The increased PC-yield ratio was observed for both donors. Sorted D7 PBs also gave rise to increased numbers of antibody secreting cells (ASCs) at D21, demonstrating that they are both phenotypically and functionally distinct from their actB counterparts (Figure 1F). We note that at D7, the cells were largely Ki-67^+^ and CD138^-^, whereas the CD138^+^ cells that emerged by D21 had substantially reduced Ki-67 levels (Figure 1F, G), consistent with their cell cycle withdrawal during the prePC-to-PC transition. We acknowledge that the D7 CD20^+^CD27^-^ actB population may contain cells with distinct intrinsic differentiation potentials, including anergic or memory B cell precursors. Nevertheless, these experiments established D7 as a critical developmental window for prePC generation, coincident with up-regulation of IRF4 and BLIMP1.

### Single-cell transcriptional profiling delineates prePCs and demonstrates correspondence of their PC progeny with BMPCs

To genomically delineate the prePC state and attempt to distinguish it from actB, PB and PC states, we performed single-cell RNA sequencing (scRNA-seq) at D7 and D19. After Leiden clustering (Figure S2A), the cells were initially consolidated into 3 cell states: actB, PB/prePC, and PC, based on top 100 upregulated DEGs (see Methods) (Figure 2A, B and Table S2). In accordance with the phenotypic analysis, D7 cells were primarily comprised of actB and PB/prePC states, whereas D19 cells were dominated by PCs (Figure 2B and S2B). Marker gene analysis supported the actB→PB/prePC→PC developmental continuum. Features of activated B-cells including GC-associated genes (e.g., *SPIB, SPI1, BCL2A1, BATF, BATF3, LTB, PTPRC, AICDA, IL2RA, IL2RG, IL4R, CCL22, CIITA, PCNA, PDCD2*, *PAX5, BID, CXCR4, CXCR5, PIM3*) were progressively downregulated along the PB/prePC→PC axis, while canonical PC genes (*JCHAIN, HERPUD1, DERL3, CD27, CD38, SSR3, SSR4, XBP1, TNFRSF17, ITM2C, MZB1, PIM2, PDIA4, PRDX4, SDF2L1*, *TXNDC5, TXNDC11*, *PRDM1*) were induced (Figure 2A; S2C; and Table S2). In the PB/prePC cluster, aside from silencing of selective B cell-genes (*HLA-DRA, HLA-DPB1, HLA-C, BID, CD81, PDCD2, SPIB, LTB, RBX1, CDK4*) and PC-gene induction (*IRF4*, *ZBTB20, CXCR4, RRBP1*), notable DEGs were associated with: cessation of B-cell activation-induced cycling or activation-induced cytidine deaminase activity (*SOX5*, *FOXP1*, *CDKN1A, CDKN1B, RB1, ATM, RAD21, RAD51B*), translational halt (*EIF2AK3* [PERK]), DNA remodeling (*HDAC9*, *MEF2A, MEF2C*, *DNMT1*, *EZH2, IKZF1, IKZF3*), ER biogenesis and the UPR (*FNDC3B*, *ATF6*, *YY1*), and calcium- or receptor-mediated signaling (*CAMK1D*, *MAPK8* (JNK1), *STAT3*, *IL21R*, *IL10RA, PIM1, PIM3*). The B cell-identity genes that remained unsilenced (*EBF1*, *PAX5*) in this cell cluster are known to also function in preventing differentiation into T or myeloid lineages. Within the PB/prePC cell cluster a subset of cells displayed decreased G1/S and G2/M cell cycle scores and decreased *MYC* expression, consistent with their exit from the cell cycle prior to a transition into the PC state (Table S3). Thus, based on the scRNAseq analysis we propose that prePCs represent the non-cycling cells within the PB/prePC scRNA-seq cluster. Accordingly, they may represent the Ki-67^-^ cells within the CD20^-^CD38^+^CD27^+^CD138^-^ compartment in our culture system.

**Figure 2.**
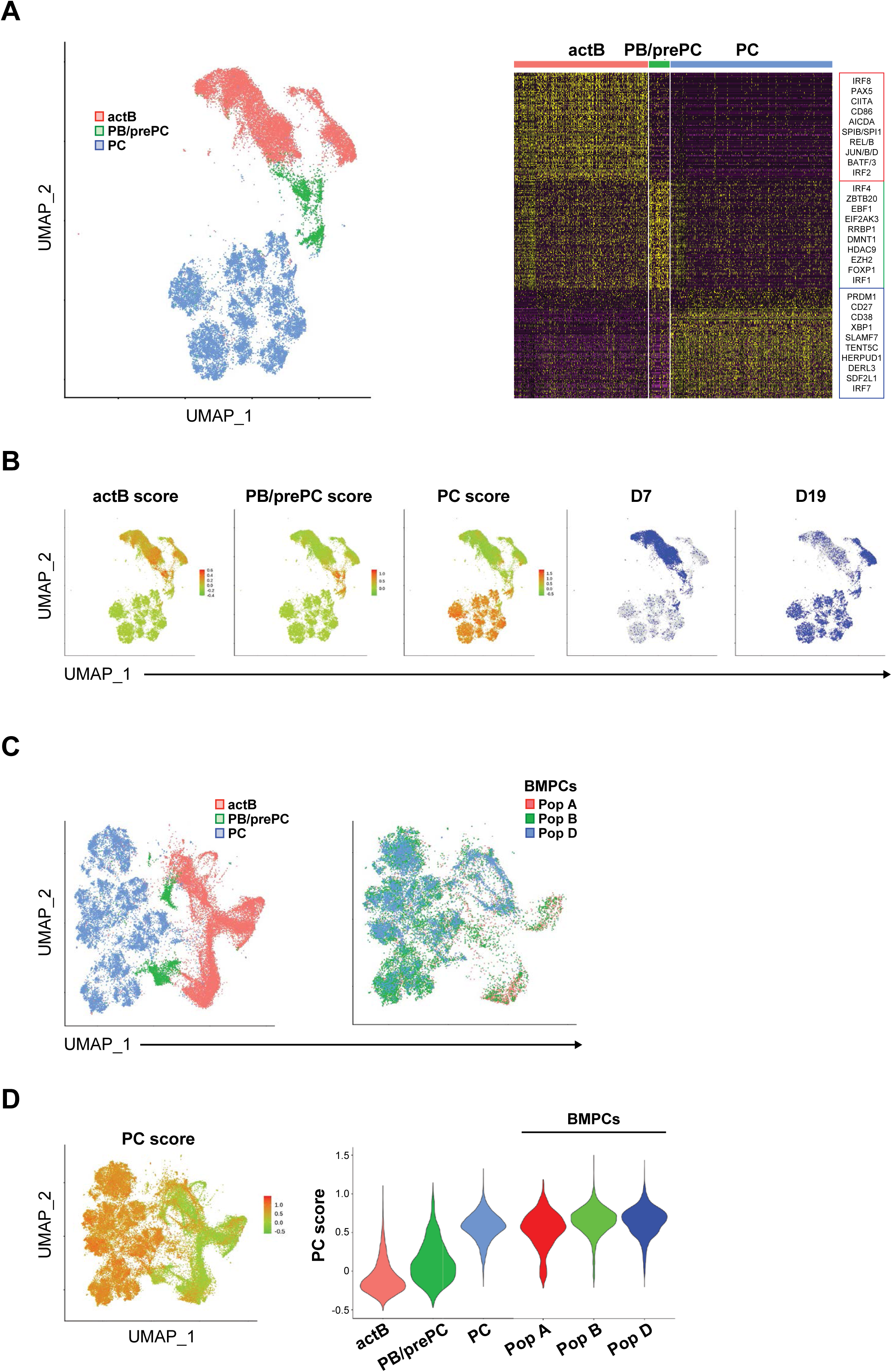
Single-cell transcriptional profiling delineates prePCs and demonstrates correspondence of their PC progeny with BMPCs. (A) *In vitro*-derived cells were profiled by scRNA-seq at D7 and D19. After Leiden clustering (Figure S2A), cells were consolidated into three major states based on marker-gene expression: actB, PB/prePC, and PC. UMAP displays aggregated cells from both timepoints and donors (Donors 1 and 2) across two independent experiments, annotated by cell state (left). Heatmap shows the top 100 marker genes per cell state, representing actB, PB/prePC, and PC gene modules (right), with representative genes from each module indicated. (B) UMAP projections show module scores for the indicated cell states (left) and timepoint annotation (right). (C) CCA-like integrated UMAPs show *in vitro*-derived cells jointly embedded with human BMPCs based on scRNA-seq profiles. (D) PC module scores, derived from the PC gene module in Figure 2A, are shown on the integrated UMAP (left) and as violin plots (right) for *in vitro*-derived cells and BMPCs grouped according to the source annotation (Pop A, Pop B, Pop D).

The PC population was largely resolved into three distinct groups of cells based on their Ig isotypes: IgM (*IGHM*^hi^), IgG (*IGHG1-4*^hi^), and IgA (*IGHA1-2*^hi^), consistent with the *in vitro* culture system being able to generate unswitched and switched PCs (Figure S2D and Table S4). *TNFRSF17* (BCMA) was broadly expressed in mature PCs, with *TNFRSF13B* (TACI) and *CCR10* expression in IgM/IgA PCs, whereas *SDC1* (CD138) transcripts were elevated in IgG PCs but did not reach significance despite ample protein expression (Figure S2D, E and Table S4), consistent with isotype-associated differences in survival signaling. Notably, the ratio of activated versus repressed DEGs shifted from 16:1 in actBs to 1:19 in PCs (Table S2), underscoring that terminal PC programming is overwhelmingly driven by gene repression.

To compare the transcriptional states of *in vitro*-generated PCs with their *in vivo* counterparts, i.e., BMPCs, we used Seurat’s SCTransform to integrate our dataset with scRNA-seq data of sorted, healthy human BMPCs (Duan et al., 2023). Integrated analysis showed that *in vitro*-derived PCs mapped closely to the *in vivo* BMPCs in low-dimensional space spanning twelve cell clusters (Figure 2C and S2F). As expected, the *in vitro* actB cells and prePCs exhibited low and intermediate PC module scores, respectively. Importantly, the *in vitro*-derived PCs exhibited high PC module scores that reached the levels of BMPCs (Figure 2D). These analyses delineate a prePC state that appears to give rise to mature PCs *in vitro* that closely resemble human BMPCs.

### Stage-specific perturbations reveal distinct functions of IRF4 and BLIMP1 in prePCs

To analyze the functions of IRF4 and BLIMP1 (*PRDM1*) within prePCs, we performed CRISPR/Cas9 editing at D7 using guide RNAs (gRNA) targeting *IRF4* (*IRF4* KO), *PRDM1* (*PRDM1* KO), or non-targeting control guide RNAs (Control) coupled with phenotypic and scRNA-seq analyses. While CRISPR/Cas9 editing was performed on bulk D7 cultures containing both activated B cells and PB/prePCs, our detailed interpretations of IRF4 and BLIMP1 functions relied primarily on downstream single-cell RNA-seq analyses of cells specifically along the prePC-to-PC trajectory. We first analyzed the impact of *IRF4* or *PRDM1* KO on PB (CD20^-^CD38^+^CD27^+^CD138^-^) and PC (CD20^-^ CD38^+^CD27^+^ICAM2^+^CD138^+^) frequencies by performing phenotypic analysis at D14 and D21 (7 or 14 days post-KO). While KO of either TF resulted in significant decreases in PB and PC frequencies, *IRF4* KO cells exhibited a more dramatic effect at the earlier developmental stage as seen by the diminished PB frequencies at 7 days post-KO (Figure 3A and S3A). Intracellular staining, 2 days post-KO, confirmed robust depletion of the targeted factor but also revealed loss of the cross-regulating TF (Figure 3B and S3B). We note that since each of the targeted TFs was largely depleted after perturbation, the impact on the targeted TF and its cross-regulated partner was assessed by analyzing the frequency of BLIMP1^hi^IRF4^hi^ cells within the total viable population. Overall, these results were suggestive of a temporally ordered model in which IRF4 acts earlier than BLIMP1. We emphasize two limits on this inference. First, editing was initiated in a heterogeneous population of actBs and PBs rather than in a purified prePC population. We therefore refer to this perturbation as stage-specific rather than prePC-restricted. Second, the bulk phenotypic data by themselves do not distinguish an earlier requirement for IRF4 from its hierarchical dominance within the coupled IRF4-BLIMP1 regulatory module (see Discussion).

**Figure 3.**
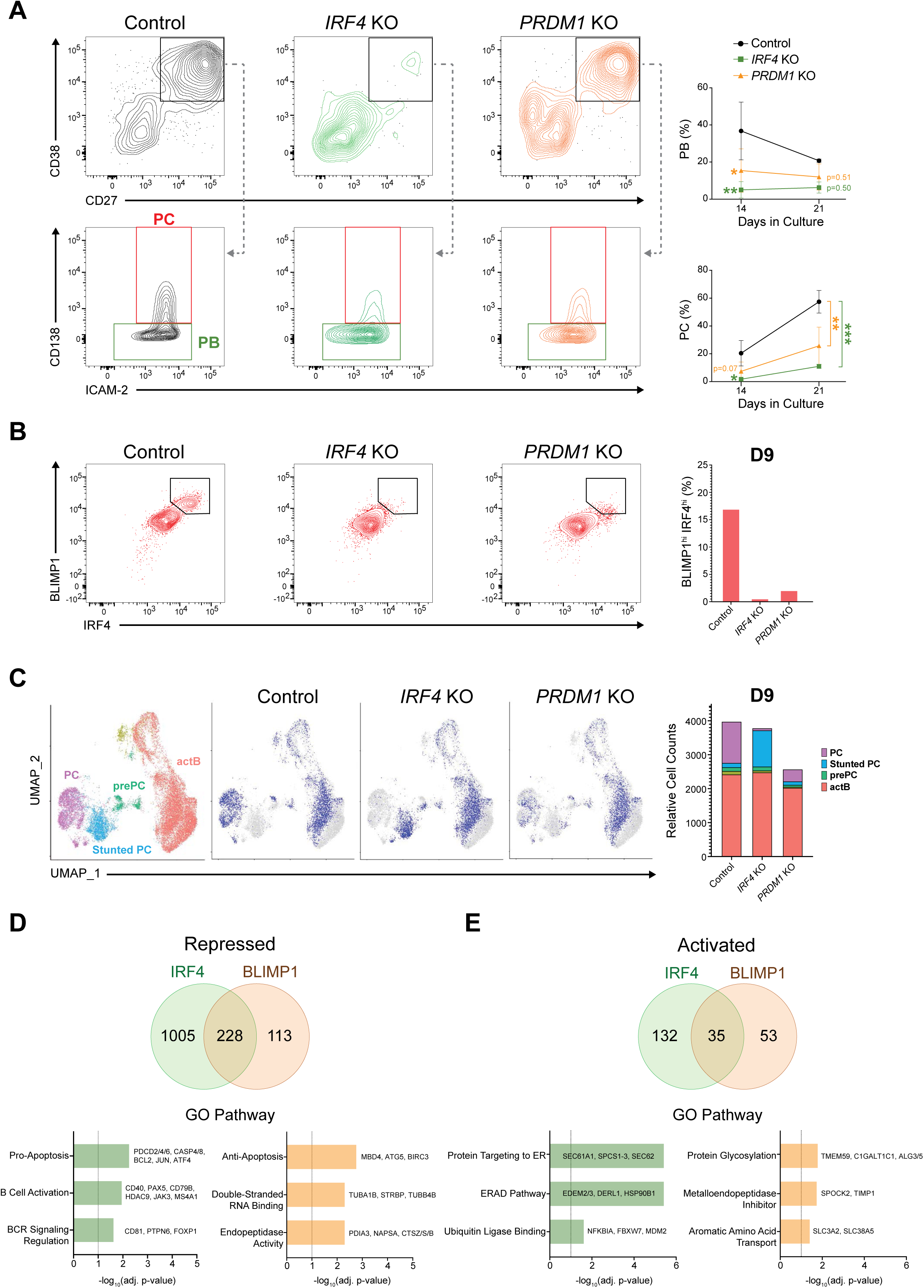
Stage-specific perturbations reveal distinct functions of IRF4 and BLIMP1 in prePCs. (A) Stage-specific *IRF4* and *PRDM1* perturbations were analyzed by immunophenotyping during PC differentiation. At D7, cells were nucleofected with CRISPR/Cas9 RNP complexes containing non-targeting control gRNAs (Control) or gRNAs targeting *IRF4* (*IRF4* KO) or *PRDM1* (*PRDM1* KO). Representative flow cytometry plots show PB (CD38^+^CD27^+^ICAM2^+/-^CD138^-^) and PC (CD38^+^CD27^+^ICAM2^+^CD138^+^) gating at D14 (left).Line graphs show mean PB and PC frequencies among live cells at D14 and D21 for Donor 1, with error bars showing S.D. (n=2-3 independent experiments; **p<0.01, ***p<0.001; two-way ANOVA with Dunnett’s post-hoc test for KO versus Control). (B) Effects of *IRF4* and *PRDM1* perturbation on IRF4 and BLIMP1 protein abundance were analyzed by intracellular flow cytometry (Donor 1 shown). Representative plots show IRF4 and BLIMP1 levels at D9, 2 days post-KO (left). Bar plot shows BLIMP1^hi^IRF4^hi^ frequencies at D9 for Donor 1 (right) (representative of 3 experiments). (C) scRNA-seq analysis of Control, *IRF4* KO, and *PRDM1* KO cells from Donor 1 at D9 from the same culture in (B). UMAP projections show aggregated cells annotated by Leiden cluster or sample identity. Displayed cells are from perturbation experiments using activated B cells from two different donors; cells shown in gray derive from the second donor, which are displayed separately in Figure S3C. Bar plot shows the distribution of cluster cell numbers across Donor 1 samples at D9 (right). (D, E) Venn diagrams show intersections of genes repressed (D) or activated (E) by IRF4 and BLIMP1, inferred from DEGs. Bar plots show selected gene ontology pathway enrichments for IRF4-only (green) and BLIMP1-only (orange) DEGs, with representative genes annotated (see Methods for DEG analysis details).

To investigate the gene expression programs that IRF4 and BLIMP1 regulate during this critical developmental window, we profiled the effects of *IRF4* or *PRDM1* KO at D9 using scRNA-seq (2 days post-KO). High-resolution scRNA-seq analyses uncovered strikingly different transcriptional consequences of perturbing *IRF4* versus *PRDM1* in prePCs (Figure 3C and S3C). Cycling PB-like cells within the PB/prePC cluster defined by the prePC module scoring were sparse at D9 in Control cells (Figure S3D and Table S5). We note that this low frequency at D9 compared to D7 (Figure 1E-G and S1F) likely reflects the difference between transcriptional cell-cycle scores and the persistence of Ki-67 protein (Table S5). While both perturbations diminished the generation of PCs, IRF4-deficient cells accumulated as an intermediate cluster that was positioned between non-dividing prePCs and PCs (Figure 3C and S3C, D). This was consistent with stunted PC maturation (blue cluster) and an earlier developmental block caused by the IRF4 perturbation that was inferred by the phenotypic analysis (Figure 3A and S3A). In contrast, a reduced fraction of BLIMP1-deficient cells progressed into a PC-like transcriptional state, suggesting that BLIMP1 is less critical than IRF4 for progression into a PC-like transcriptional state, but remains essential for the efficient consolidation of the mature PC program (Figure 3C; S3C; and Table S5). We note that Figure 3B and 3C represent distinct readouts: flow cytometry directly measures IRF4 and BLIMP1 protein abundance, whereas scRNA-seq resolves the higher resolution transcriptional states of the persisting cells after perturbation despite continued transcript presence. As a particular example, *IRF4* and *PRDM1* KOs led to a failure to repress *ID3* in the Stunted PC cluster and *PRDM1* KO PCs, respectively (Figure S3D). The combined phenotypic and genomic analyses were consistent with a model involving temporally ordered regulatory actions of IRF4 and BLIMP1 in PC precursors (see Discussion).

To analyze the transcriptional programs regulated by IRF4 and BLIMP1 during the prePC-to-PC transition, we performed differential gene expression analyses on the two most pronounced clusters of cells impacted by each perturbation along the PC trajectory: *IRF4* KO-dominated “Stunted PCs” versus all “PCs” and “*PRDM1* KO” PCs versus “Control” PCs (Figure 3D, E; S3E-J; and Table S6). As expected, changes in gene expression levels were highly correlated with changes in the frequency of expressing cells for both perturbations (R^2^_IRF4_=0.8732, R^2^_BLIMP1_=0.8423) and revealed 1,400 significant *IRF4* KO differentially expressed genes (DEGs) (167 down-regulated and 1,233 up-regulated) and 429 significant *PRDM1* KO DEGs (88 down-regulated and 341 up-regulated) (Figure S3E). Genes regulated discordantly by the two perturbations were analyzed separately (Figure S3F) and are excluded from the concordant intersections reported below in Figure S3G, H. Consistent with the repression-dominated logic of PC programming (Figure 2A), *IRF4* KO DEGs showed a 7:1 ratio of up-regulated to down-regulated genes, and *PRDM1* KO DEGs a 4:1 ratio, demonstrating that both TFs dominantly function as repressors during the prePC-to-PC transition (Figure S3F).

Intersecting these DEG sets (Figure 3D, E and S3F-J) revealed both convergent and divergent genomic programming by IRF4 and BLIMP1 during the prePC-to-PC transition. The 228 genes that were co-repressed by IRF4 and BLIMP1 were highly enriched for SPI1 (PU.1) target genes based on ChIP-X Enrichment Analysis (ChEA), as well as cadherin and ubiquitin ligase binding pathways (GO; Figure S3G), while the 35 co-activated genes were highly enriched for endoplasmic reticulum (ER) unfolded protein response (UPR) and ER-associated protein degradation (ERAD) pathways (GO; Figure S3H). IRF4-only repressed genes (1,005) were enriched for MYC target genes (ChEA; Figure S3I) as well as pro-apoptosis, B-cell activation/proliferation, BCR-signaling, and NF-κB-signaling pathways (GO; Figure 3D). In contrast, BLIMP1-only repressed genes (113) were uniquely enriched for putative ZBTB7A target genes (ChEA; Figure S3I) and anti-apoptosis pathways specifically utilized by actB for activation-induced survival (GO; Figure 3D). IRF4-only activated genes (132) were uniquely enriched for NFYA/NFYB target genes (ChEA; Figure S3J) and ER pathways (GO; Figure 3E), while BLIMP1-only activated genes (53) were uniquely enriched for NELFE target genes (ChEA; Figure S3J) and the protein glycosylation pathway (GO; Figure 3E). The complete ChEA and GO pathway analyses can be found in Table S7. Together, these results support a model in which IRF4 independently initiates silencing of B cell activation, proliferation, and BCR signaling programs while inducing *PRDM1* expression and priming ER pathway genes. BLIMP1 subsequently extinguishes activated B cell survival genes and, together with IRF4, co-activates a selective set of ER/UPR genes that support antibody secretion, reflecting a coupled, temporally ordered execution of the mature PC transcriptional program.

### Differential IRF4 and BLIMP1 genomic occupancy reveals a discriminating ISRE/EICE motif lexicon

The divergent transcriptional phenotypes prompted us to ask whether IRF4 and BLIMP1 engage distinct as well as shared *cis*-regulatory landscapes during PC differentiation. We performed CUT&RUN assays for IRF4 and BLIMP1 at the prePC developmental window (D7) and intersected high-confidence genomic binding regions (CUT&RUN peaks) with multiome-defined open chromatin regions (OCRs) (Figure 4A). The IRF4/BLIMP1 bound OCRs were then associated with KO DEGs (within 200kb of transcription start sites) and scanned for cognate binding motifs. To systematically compare the genes and motifs associated with genome-wide IRF4 and BLIMP1 binding patterns, we first stratified all OCRs into three categories: IRF4-only bound (IRF4-bound), BLIMP1-only bound (BLIMP1-bound), and overlapping (co-bound) OCRs (Figure 4A). Notably, a substantial fraction of IRF4 and BLIMP1 binding regions overlapped when using all OCRs as the background set (hypergeometric p≈0), consistent with considerable co-occupancy of accessible regulatory elements during the prePC-to-PC transition as previously reported in murine B cells (Minnich et al., 2016). As anticipated, IRF4 and BLIMP1 related motifs (EICE, ISRE, and BLIMP1) were enriched within each OCR class (Figure 4B and S4A).

**Figure 4.**
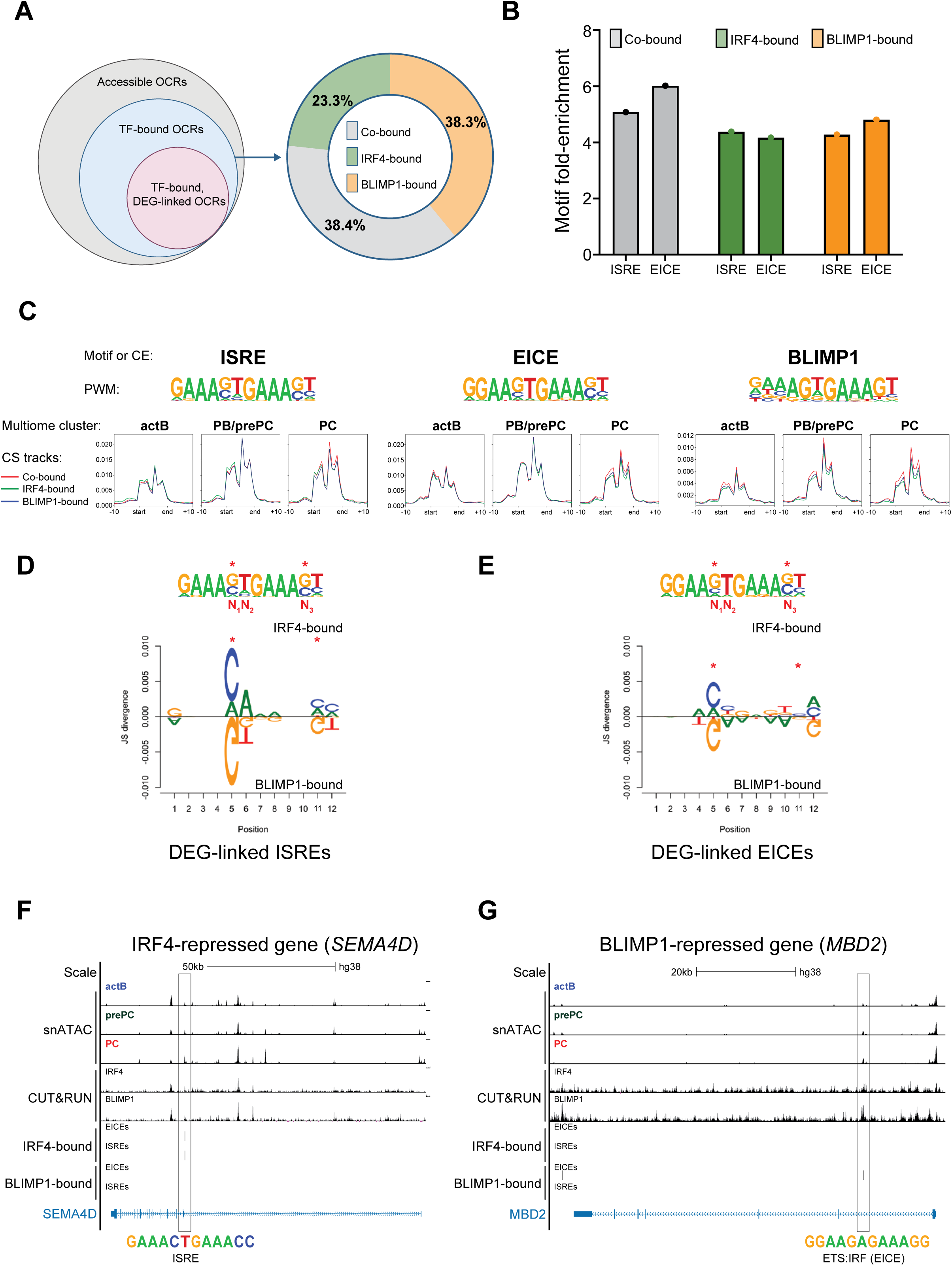
Differential IRF4 and BLIMP1 genomic occupancy reveals a discriminating ISRE/EICE motif lexicon. (A) Schematic summarizes the strategy for comparing IRF4- and BLIMP1-bound open chromatin regions (OCRs) linked to KO DEGs (left). Donut chart shows the distribution of IRF4-bound, BLIMP1-bound, and co-bound OCRs (p≈0 for overlapping region frequency, hypergeometric test) (right). (B) Bar plot shows enrichment of ISRE and EICE motifs in co-bound (gray), IRF4-bound (green), and BLIMP1-bound (orange) OCRs (FDR≤0.056). (C) State-specific ChromBPNet contribution-score analysis of ISRE, EICE, and BLIMP1 motif instances across actB, PB/prePC, and PC states. PWMs for the indicated motifs are displayed above histograms showing contribution-score signal at motif instances in the indicated OCR categories. (D, E) Sequence logos show Jensen-Shannon (J-S) divergence between (D) ISRE or (E) EICE motif instances at IRF4-bound OCRs linked to *IRF4* KO DEGs and BLIMP1-bound OCRs linked to *PRDM1* KO DEGs. Reference motifs above divergence logos indicate degenerate nucleotides of interest (red asterisks), along with nomenclature for variant positions. Discriminating variant positions in J-S divergence logos are denoted by red asterisks. (F) UCSC Genome Browser tracks show an example of an IRF4-bound ISRE linked to the IRF4-repressed gene *SEMA4D*. (G) UCSC Genome Browser tracks show an example of a BLIMP1-bound EICE linked to the BLIMP1-repressed gene *MBD2*.

To relate genomic occupancy with base-pair-level regulatory logic, we used ChromBPNet modeling to infer the contribution of each base-pair to chromatin accessibility across ISRE/EICE/BLIMP1 motif instances within each peak set (co-bound, IRF4-only, BLIMP1-only bound) and cell state (actB, PB/prePC, PC). State-specific ChromBPNet models revealed that the contribution of ISRE/EICE/BLIMP1 motifs to chromatin accessibility increased considerably between the actB and PB/prePC states and remained elevated in PCs (Figure 4C). Notably, ISRE/EICE/BLIMP1 motifs at each peak set category exhibited similar increases in their contribution scores in PB/prePC and PC states, consistent with coordinated increases in IRF4 and BLIMP1 regulatory activity at these elements during the prePC-to-PC transition. However, this also highlighted the paradox in accounting for the distinct genomic actions of IRF4 and BLIMP1 in prePCs.

Given that ISRE/EICE motifs were enriched in IRF4-only, BLIMP1-only, and co-bound OCRs (Figure 4B), motif class alone could not explain the distinct transcriptional effects of *IRF4* and *PRDM1* perturbations. We therefore asked whether nucleotide variation within each motif class discriminates IRF4 from BLIMP1 occupancy at selected regulatory regions. To do so, we filtered all IRF4- and BLIMP1-bound OCRs associated with DEGs (±200kb of TSS), and regenerated TF-specific ISRE and EICE PWMs to calculate their Jensen-Shannon (J-S) divergence, a measure of nucleotide bias, plotted as logos (Figure 4D, E and S4B, C). This comparative PWM analysis revealed nucleotide preferences at 5’ (N_1_/N_2_) and 3’ (N_3_) positions flanking the 3’-GAAA half site (N_1_N_2_-GAAA-N_3_) that differed between IRF4- and BLIMP1-bound regions. BLIMP1-biased ISREs or EICEs enriched for G_1_T_2_-GAAA-G_3_ and G_1_A_2_-GAAA-G_3_ sequences (Figure 4D, E). In contrast, IRF4-bound ISREs or EICEs were biased for distinct flanking nucleotides: (C/A)_1_A_2_-GAAA-(C/A)_3_ and (C/A)_1_(C/T)_2_-GAAA-(C/A)_3_. These PWM differences suggest that motif degeneracy, particularly at N_1_ (ISRE/EICE) and N_3_ (ISRE) positions, can encode specificity that could bias TF binding, thereby enabling distinct genomic actions of IRF4 and BLIMP1 on ISREs and EICEs. For example, *SEMA4D*, which encodes for a transmembrane protein that promotes SHP1 dissociation from CD72 (Kumanogoh et al., 2000; M. et al., 2017), is uniquely repressed by IRF4 (Figure 3D and Table S2) and linked to an IRF4-bound ISRE that contains an IRF4-biased sequence (C_1_T_2_-GAAA-C_3_) (Figure 4F). In contrast, *MBD2*, which encodes a DNA methylation binding protein that indirectly promotes BCR signaling (Jing et al., 2025), is uniquely repressed by BLIMP1 and linked to a BLIMP1-bound EICE that contains a BLIMP1-biased sequence (G_1_A_2_-GAAA-G_3_) (Figure 4G). Similar TF-specific, activated DEGs supporting Ig secretion (*XBP1*; (Shaffer et al., 2004)) and PC survival (Defender against death 1, *DAD1*; (Nguyen et al., 2025)) were associated with cognate TF-biased motifs (Figure S4D, E). Together, these data suggested a model in which IRF4 and BLIMP1 frequently converge on related IRF-like motifs but exhibit distinct occupancy on a subset of ISRE- and EICE-containing regions, dictated by their base-pair preferences at the same two degenerate positions within both motifs. We note that the regenerated PWMs identify both C and A as IRF4-biased at N_1_/N_3_, whereas the EMSAs described below show that IRF4 alone does not detectably bind A-containing ISRE variants. The EMSA-derived IRF4-specific category used in the analyses that follow is therefore the data-supported C-containing subset.

### *In vitro* TF binding assays test base-pair rules and predict genomic partitioning of IRF4 and BLIMP1

To directly test the functional impact of motif degeneracy on TF binding, we performed EMSAs using *in vitro*-translated proteins and a comprehensive panel of DNA probes with single nucleotide variants derived from a shared EICE-embedded sequence in the *CIITA* promoter (GGAA-N_1_T_2_-GAAA-N_3_). Notably for these single base-pair permuted DNA probes, the N_2_ position was maintained as an invariant T given its importance in the BLIMP1 core and prevalence in the CUT&RUN bound regions for both TFs across ISREs and EICEs. For EICE variant probes, PU.1::IRF4 heterodimers and IRF1, a prototypical IRF, had relatively relaxed specificities capable of binding S/N or N/S (12/16 probes, PU.1:IRF4), and N/N (16/16 probes, IRF1), respectively (Figure 5A). Note that IUPAC codes are used for the N_1_/N_3_ pair with S = C or G; W = A or T; H = A, C or T; N = any nucleotide. In contrast with the above, BLIMP1 binding was restricted to G/N or N/G variants (7/16 probes), consistent with the observed J-S divergence (Figure 4D, E). To investigate whether similar binding preferences exist for ISREs, we converted the same EICE-embedded sequence to an ISRE by replacing the guanine at position 2 of the first IRF half-site (GGAA to GAAA), yielding GAAA-N_1_N_2_-GAAA-N_3_. Interestingly, while BLIMP1 exhibited indistinguishable ISRE-binding preferences (G/N or N/G) compared to its binding of EICE probes, IRF4 alone displayed markedly more restrictive binding to ISRE variants (S/S only, 4/16 probes), contrasting sharply with its cooperative binding mode with PU.1 to EICE probes (Figure 5B). We note that IRF4 did not display enhanced binding with PU.1 to ISRE probes (Figure S5A) confirming that ETS-cooperative binding is strictly EICE-dependent. In contrast to IRF4 and BLIMP1, IRF1 detectably bound all variants tested (N/N), consistent with its broader recognition of ISRE sequences (16/16 probes, Figure 5A, B). Thus, a clear hierarchy of nucleotide selectivity emerged: IRF4 is most restrictive at ISREs, followed by BLIMP1 at both EICEs and ISREs, then PU.1::IRF4 at EICEs, and finally IRF1 at both motif types.

**Figure 5.**
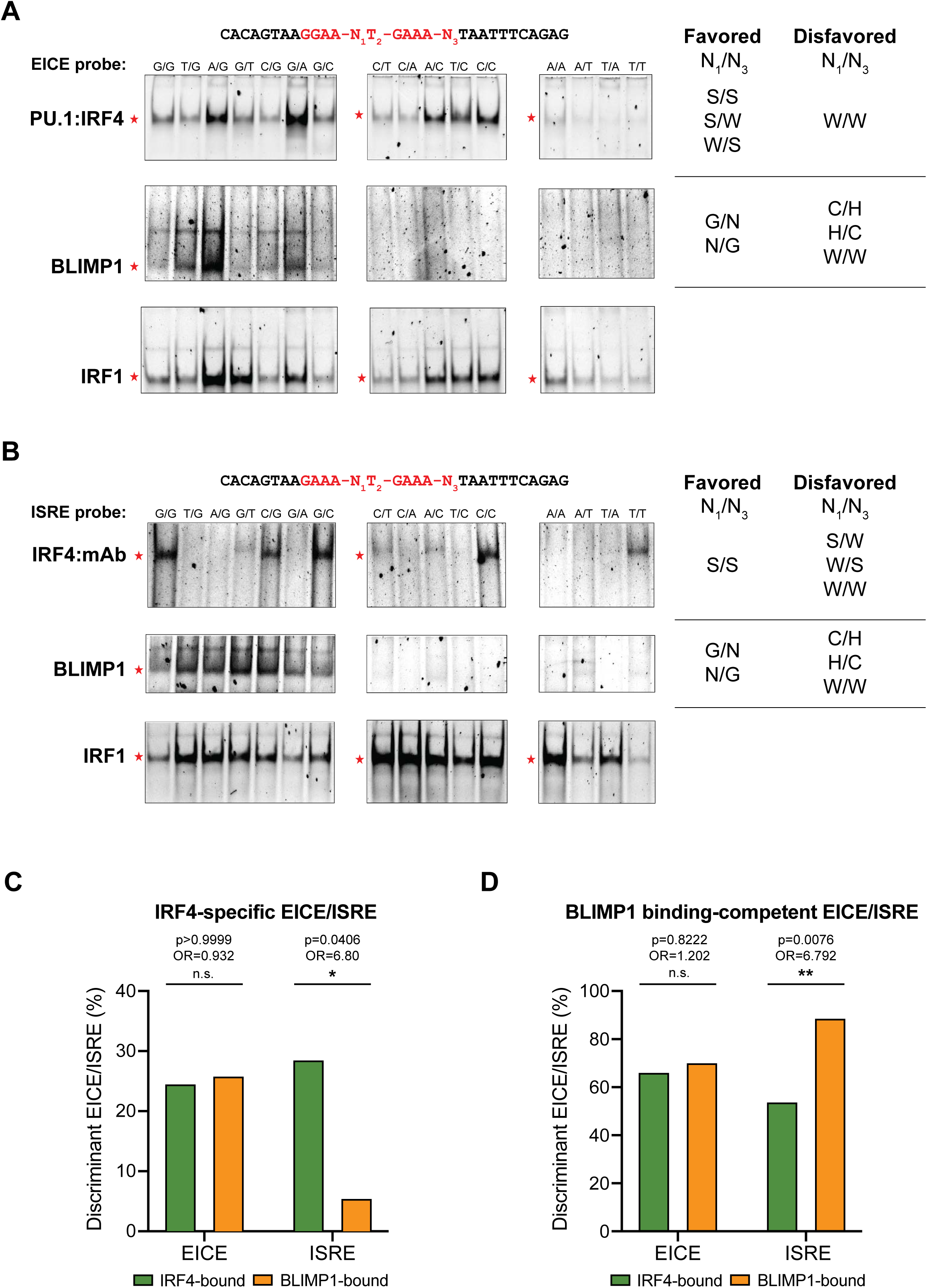
*In vitro* TF binding assays test base-pair rules and substantiate the ISRE/EICE regulatory logic. (A, B) Electrophoretic mobility shift assays (EMSAs) were performed to test N_1_/N_3_ base-pair rules for IRF4 and BLIMP1 binding at EICE and ISRE motif variants. Gels are grouped by G-containing (G/N or N/G), C-containing without G (C/H or H/C), and A- or T-containing (W/W) variants (N_1_/N_3_). Relevant TF-specific bands are annotated with red stars. Summaries of favored and disfavored N_1_/N_3_ nucleotides for each TF-probe combination are shown on the right. (A) Cell-free *in vitro*-transcribed/translated (IVT) lysates containing IRF4 and PU.1 (top), BLIMP1 (middle), or IRF1 (bottom) were assayed for binding to labeled EICE probes. (B) IVT lysates containing IRF4 (top), BLIMP1 (middle), or IRF1 (bottom) were assayed for ISRE variant probe binding. IRF4-ISRE assays were performed with an antibody directed against the epitope-tag and analyzed for super shifted complexes. (C, D) Statistical enrichment analysis of discriminating N_1_/N_3_ motif variants at IRF4-bound OCRs linked to IRF4 DEGs (green) or BLIMP1-bound OCRs linked to BLIMP1 DEGs (orange). Bar plots show the frequency of (C) IRF4-specific ISRE or EICE motif variants (N_1_/N_3_=C/C) and (D) BLIMP1 binding-competent ISRE or EICE motif variants (N_1_/N_3_=G/N or N/G) (Fisher’s exact tests; see Table S8 for counts).

We next asked whether this biochemically defined binding hierarchy is reflected in the partitioning of TF genomic occupancy. We note that OCRs associated with *IRF4* or *PRDM1* KO DEG sets were not differentially enriched for ISRE-versus EICE-motifs (Figure S5B). Thus, preferential action of the TFs at one of the two motif classes was deemed unlikely to account for the differential genomic activity of IRF4 and BLIMP1. We instead hypothesized that differential motif lexicon within each motif class is responsible for directing IRF4 versus BLIMP1 regulatory activities. To test this hypothesis, we used the EMSA-derived binding categories (“Favored” variants from Figure 5A, B), and re-binned ISREs and EICEs within IRF4-bound DEG-linked OCRs and BLIMP1-bound DEG-linked OCRs. The results revealed a significant partitioning at ISREs: IRF4-specific ISREs (GAAA-C_1_N_2_-GAAA-C_3_) showed 6.8-fold higher odds of occurring at IRF4-bound DEG-linked OCRs compared to BLIMP1-bound DEG-linked OCRs (OR=6.80, Fisher’s p=0.0406), and reciprocally, BLIMP1 binding-competent ISREs (GAAA-[G/N]_1_N_2_-GAAA- [N/G]_3_) showed 6.8-fold higher odds of occurring at BLIMP1-bound DEG-linked OCRs compared to IRF4-bound DEG-linked OCRs (OR=6.79, Fisher’s p=0.0076) (Figure 5C, D). Contingency tables are provided in Table S8. Both tests are two-sided Fisher’s exact tests; between the two tests, the BLIMP1-directed ISRE comparison remained significant after Bonferroni correction (p_corr_ threshold≤0.025) whereas the IRF4-directed comparison lost significance by a marginal amount. These odds ratios provide quantitative support for the motif lexicon model: N_1_/N_3_ nucleotide identity within ISREs predicts preferential association with IRF4-versus BLIMP1-bound DEG-linked OCRs. In contrast, IRF4-specific and BLIMP1 binding-competent EICEs showed slight biases toward BLIMP1-binding regardless of binning, but did not reach statistical significance (Figure 5C, D), consistent with EICEs being more promiscuous for both factors. We ascribe this observation to cooperative IRF4::PU.1 binding at EICEs, with PU.1 providing additional binding energy that partially overrides IRF4-intrinsic N_1_/N_3_ nucleotide preferences and reduces the discriminating power of variant nucleotides at EICEs *in vivo*. Together, the biochemical and genomic analyses support a differential motif lexicon model in which N_1_/N_3_ nucleotide identity within ISREs, and to a lesser extent EICEs, discriminates IRF4 from BLIMP1 binding and encodes the transition from the IRF4-promoted initiation to BLIMP1-assisted consolidation of the human PC transcriptional program.

## Discussion

This study defines a stage-specific IRF4–BLIMP1 regulatory logic for human PC differentiation and delineates an ISRE/EICE motif lexicon as a mechanism that partitions their genomic activities. By combining a stepwise human B cell differentiation system with stage-specific perturbation at D7, single-cell transcriptomics, chromatin profiling, CUT&RUN, and biochemical DNA-binding assays, we show that IRF4 acts early to license the prePC-to-PC transition, whereas BLIMP1 consolidates the mature secretory state. The system further enables characterization of transient human prePCs, which have been difficult to study directly, and shows that D7 CD20^-^CD27^hi^CD38^hi^CD138^-^ PBs are enriched for precursors that generate transcriptionally BMPC-like PCs. A related study has profiled the activation and differentiation of human B cells into ASCs (up to D11) by scRNA-seq (Verstegen et al., 2023). However, this study did not analyze the generation of CD138^+^ PCs or utilize APRIL, TGF-β3 or IL-6 in their culture conditions.

We have previously used adoptive transfer studies coupled with scRNA-seq analyses to characterize murine PC precursors emanating from germinal centers that give rise to bone marrow plasma cells (Manakkat Vijay et al. 2024). Related work in humans has defined circulating CD138^-^ ASCs after vaccination that may represent PC precursors (Ferreira-Gomes et al., 2024). In our human *in vitro* system, prePCs are enriched within CD20^-^CD27^hi^CD38^hi^CD138^-^ PBs at day 7, aligning with ex vivo observations that antigen-induced circulating plasmablasts peak around day 7 post-immunization. The emergence of *CCR10*^+^ and *CXCR3*^+^ PCs in the *in vitro* system and their near absence in *IRF4* and *PRDM1* KO cultures suggests that parts of the PC subset differentiation program are intrinsically coupled to terminal differentiation rather than requiring initial niche imprinting or priming. In agreement with recent single cell profiling of primary tonsillar human B cells which described a CD30⁺ (*TNFRSF8*) subset representing a prePC state (Fields et al., 2026), we found that *TNFRSF8* transcripts are expressed in actBs, increase in amplitude in PB/prePCs, and are extinguished in PCs. This system should enable genetic and chemical perturbation screens for regulators of human PC differentiation and survival (D’Souza et al., 2026; Nguyen et al., 2018) and future optimization of BM-like survival conditions or humanized-mouse engraftment assays could further test the developmental competence of *in vitro*-generated prePCs (Cheng et al., 2022; Hill et al., 2026; Nguyen et al., 2018).

IRF4 and BLIMP1 participate in early B cell activation programs, GC-associated states, and terminal differentiation; perturbations induced at earlier stages cannot therefore distinguish their requirements for initiating ASC commitment from those needed to promote PC maturation and maintenance. By editing *IRF4* or *PRDM1* at the D7 prePC-generating window, we demonstrate that both TFs remain essential along the PC trajectory, consistent with murine genetic evidence and regulatory models in which graded IRF4 accumulation drives *PRDM1* induction and PC commitment. Notably, loss of IRF4^hi^ BLIMP1^hi^ cells after either KO supports a model of mutual reinforcement and positive feedback during PC differentiation. Analyses of the perturbed prePC states by scRNA-seq reveal that PC maturation is overwhelmingly dominated by gene repression: the number of silenced genes exceeds the set of induced genes in both perturbations, with *IRF4* KO DEGs showing a 7:1 repression-to-activation ratio and *PRDM1* KO DEGs a 4:1 ratio. This asymmetry reinforces a distinctive feature of PC programming: the mature PC state is comparatively compact and specialized, genomically focused on sustaining high immunoglobulin synthesis and secretion via proteostasis and UPR elaboration under chronic ER stress. In contrast, the activated B cell state comprises broader expression modules that enable antigen sensing, signaling responsiveness, proliferation, immunomodulation, and chemotaxis (Rastogi et al., 2022). The scale of genome-wide repression by IRF4 and BLIMP1 suggests that sustained expression of B cell genes unrelated to secretion is incompatible with terminal maturation, consistent with a model in which repression of B cell gene regulatory networks is a prerequisite for the emergence and maintenance of the PC secretory state.

Despite the shared phenotypic consequence, i.e., strong impairment of PC generation, after loss of either TF, the IRF4 perturbation yielded a distinctive stunted state that reveals a non-redundant circuitry within the IRF4-BLIMP1 module. Although *IRF4* and *PRDM1* perturbations each reduce expression of the other factor within two days, the distinct *IRF4* KO stunted intermediate indicates that the initiating perturbation and the resulting kinetics of TF decay are not functionally equivalent. A parsimonious interpretation suggests that IRF4 contributes an early "licensing" function through repression of B cell identity genes and priming of secretory and proteostasis programs, acting independently of BLIMP1. Our findings of temporally ordered action of IRF4 and BLIMP1 in human PC precursors could explain the failure of activated murine B cells to differentiate into PCs under signaling conditions that induce BLIMP1 but not IRF4 (Luo et al., 2023). The licensing functions of IRF4 could include repression of BCL6-driven programs through induction of the antagonistic factor *ZBTB20*, alongside repression of *IRF8* and *ID3*, to facilitate GC exit and prePC commitment (Chevrier et al., 2014; Gloury et al., 2016). Dysregulation of chemotactic receptors in the *IRF4* KO stunted state, including elevated *CXCR4* and *CD44* expression, suggests that IRF4 also participates in rewiring cell migratory properties during differentiation, a function that BLIMP1 cannot compensate. This model fits with the temporal logic in which IRF4 induction precedes and promotes *PRDM1* expression, providing a window for IRF4 to act in advance of accumulating BLIMP1 protein. IRF4 and BLIMP1 therefore operate as coupled TFs whose functions appear temporally ordered and subsequently concerted across the prePC-to-PC transition. Thus, our data suggest IRF4 initiates critical regulatory transitions while BLIMP1 consolidates terminal PC fate as previously shown via reinforcing repression of the B cell program, and stabilizing survival and secretory gene modules (Conter et al., 2025; Tellier et al., 2016). While our data support a temporally ordered model in which IRF4 acts early to license the prePC-to-PC transition and BLIMP1 consolidates the terminal state, these experiments do not exclude alternative explanations. Because both factors are mutually reinforcing, the observed phenotypic divergence between *IRF4* and *PRDM1* KO could also reflect differences in editing efficiency, protein stability and decay kinetics, or hierarchical dominance. Future studies utilizing degron-based perturbations, rescue/add-back experiments, and gain-of-function analyses together with high-resolution time courses of IRF4 and BLIMP1 protein clearance after each perturbation, will be needed to resolve their functional ordering. If the principal role of IRF4 at this stage is to induce *PRDM1*, ectopic BLIMP1 expression in *IRF4*-deficient prePCs should rescue the stunted phenotype; failure to rescue would establish an independent early licensing function for IRF4.

The co-repressed gene set (228 genes enriched for SPI1/PU.1 targets) suggests a further mechanistic distinction between IRF4 and BLIMP1 in dismantling B cell identity. IRF4 may act competitively at the protein level by displacing IRF8 from ETS:IRF complexes at EICEs, thereby remodeling the transcriptome during the early prePC window. This in turn implies further molecular differences in the transcriptional regulatory domains of IRF4 and IRF8. BLIMP1 instead acts at the transcriptional level by repressing *SpiB* and *SPI1* (PU.1) expression, thereby blocking the maintenance of ETS:IRF complexes. Thus, both TFs appear to ultimately inhibit ETS:IRF-driven B cell identity gene expression in prePCs, but through mechanistically distinct and temporally ordered strategies, protein-level displacement by IRF4 followed by transcriptional elimination of the ETS partners by BLIMP1.

By integrating *IRF4*- and *PRDM1*-perturbed prePC states with multiome and CUT&RUN profiling, we uncover a predictive motif lexicon that may be used to partition the distinct versus coordinated genomic actions of the two PC fate determinants. ChromBPNet modeling supports a model in which IRF4 acts competitively at ISREs by displacing activating prototypical IRFs, whereas BLIMP1 may achieve more durable repression by displacing all bound factors at its preferred G/N or N/G EICE/ISRE variants. These results are consistent with competition EMSAs showing displacement by BLIMP1 of PU.1:IRF4 complexes bound to G/G-variant EICEs (Minnich et al., 2016). Multiple non-mutually exclusive mechanisms likely additionally contribute to these effects, including:(i) direct repression via recruitment of corepressor and chromatin-modifying complexes at B cell identity loci; (ii) feed-forward activation of secondary TFs and stress-adaptation regulators that amplify secretory capacity; and (iii) chromatin gating, in which one TF establishes or stabilizes regulatory element accessibility to enable productive engagement by the other. Our motif analyses refine these possibilities by revealing that nucleotide-level lexicon within ISRE and EICE elements predicts differential versus overlapping binding, providing a DNA sequence-grounded rationale for how IRF4 and BLIMP1, despite recognizing related motif sequences within a tightly coupled positive-feedback architecture, could partition their genomic targets and execute ordered functions along the prePC-to-PC trajectory. While the EMSA and CUT&RUN as well as multiome analyses support the motif lexicon as a predictive model for differential TF occupancy, direct causal testing of endogenous regulatory elements, for example, by base editing of selected ISRE/EICE variants in differentiating cells, will be required to determine if these nucleotide variants are sufficient to predictably alter gene activity.

Our results suggest that ISRE/EICE motif lexicon may reflect a general principle by which *cis*-regulatory micro-variation tunes the outputs of conserved IRF/BLIMP1-controlled gene circuits across immune contexts. IRF family TFs are repeatedly deployed across myeloid and lymphoid lineages to control activation, effector functions, and differentiation, yet closely related factors yield distinct cell-state outputs (Glasmacher et al., 2012; Honda & Taniguchi, 2006). A potential precedent for the temporally ordered IRF4-then-BLIMP1 logic exists in CD4/8 T cell effector differentiation, with a parallel to PC fate determination. IRF4 is required for T_h_2, T_h_9, T_h_17, and T_FH_ differentiation, as well as T_reg_ effector function; in CD8 T_eff_, IRF4 is required for cytotoxic effector or T_c_9/T_c_17 differentiation and memory formation (Huber & Lohoff, 2014). In both T and B cell contexts, TCR or BCR signal strength drives graded IRF4 accumulation that acts as a fate switch, promoting terminal effector or PC commitment over memory or GC fates, respectively (Cretney et al., 2011; Harberts et al., 2021; Jain et al., 2016; Lohoff et al., 2002; Nayar et al., 2014; Schmidt et al., 2023; Sciammas et al., 2006; Tominaga et al., 2003; Yao et al., 2013; Zou et al., 2020). In both lineages, IRF4 likely displaces the antagonizing IRFs like IRF8, and promotes induction of the *PRDM1* gene. BLIMP1 then consolidates the terminal fate by repressing an overlapping set of alternative-fate transcription factors: *BCL6*, *TCF7*, *ID3*, and *IRF8* in CD8 T_eff_; *BCL6*, *PAX5*, *EBF1*, *ID3*, and *IRF8* in PCs, while driving metabolic and functional reprogramming suited to high-output cytotoxic or secretory activity. The *IRF4* KO phenotype in CD8 T cells mirrors that observed here in PCs: an impaired consolidation of the terminal program and residual expression of alternative-fate genes (Nayar et al., 2014; Zou et al., 2020). The suggestion that the IRF4-BLIMP1 module may use related cis-regulatory logic to generate terminal effector states across lymphocyte lineages remains speculative and will require directed experimental testing.

## Materials and Methods

### Tissue Culture Media

RPMI-1640 (Gibco, 61870-036) was supplemented with 10% (v/v) heat-inactivated (HI) FBS (Cytiva, SH30071.03), 55µM ß-mercaptoethanol (ß-ME; Sigma-Aldrich, M6250-250ML), 1% HEPES (Gibco, 15630-080), 100U/mL penicillin/streptomycin (P/S; Gibco, 15140-122), 1x MEM non-essential amino acids (NEAA; Gibco, 11140-050), 1mM sodium pyruvate (Gibco, 11360-070), and 0.5x GlutaMAX (Gibco, 35050-061). IMDM (Gibco, 12440-053) was supplemented with 10% HI FBS, 55µM ß-ME, 1% HEPES, 100U/mL P/S, and 1x NEAA. FACS buffer is DPBS without Mg^2+^/Ca^2+^ (Gibco, 14190-144) supplemented with 3% HI FBS and 2mM EDTA mixed and passed through a 0.22µm sterile filter.

### Mitomycin C treatment of MS40L^lo^ (CD40L-expressing) cells

Stock mitomycin C prepared in DMSO (1mg/mL; Sigma-Aldrich, M4287-2MG) was diluted to 10µg/mL (1:100) in complete IMDM media immediately prior to resuspending and plating 2x10^4^ MS40L^lo^ cells in 6-well TC culture plates (Corning, 07-200-83) for mitotic arrest. Cells were incubated for 6h and wells were washed 3x with DPBS before overnight recovery in complete IMDM media. IMDM was replaced with complete RPMI-1640 media the next day before the addition of naïve B cells.

### Naïve B cell isolation and PC culture system

Naïve B cells were isolated from commercial healthy-donor PBMCs (STEMCELL Technologies) by negative-selection bead-enrichment (STEMCELL Technologies, 17254). D0-4 naïve B cells were co-cultured with 2x10^4^ mitomycin C-treated MS40L^lo^ cells (gift from Dr. Garnett Kelsoe), supplemented with BAFF (BioLegend, 559604), IL-4 (BioLegend, 574004), IL-2 (BioLegend, 589104), and IL-21 (Gibco, 20021100UG) as previously described (Su et al., 2016). Subsequent culture steps followed those described by the Tooze laboratory (Cocco et al., 2012). Briefly, D4 activated B cells were pooled for each donor, then filtered twice to remove MS40L^lo^ cells. Filtered cells were cultured in IL-2, IL-21, APRIL (added on Day 5, Gibco, 31010C100UG), and chemically defined lipids (CDLs, Gibco, 11905031) for D4-7. D7 plasmablasts were filtered once for residual MS40L^lo^ cells and then cultured in IL-6 (Gibco, 20006100UG), IL-21, APRIL, TGF-ß3 (Gibco, 10036E100UG), and CDLs for D7-14, with media replacement on D11 for cultures going longer than D11. D14+ cells were cultured in IL-6, APRIL, TGF-ß3, and CDLs with media replacement every 3-4 days as necessary.

### CRISPR-Cas9/RNP-editing

Using P3 buffer (Lonza Amaxa, V4XP-3024) and program CA-137 for the 4D-Nucleofector X Unit (Lonza Amaxa, AAF-1003B/X) per the manufacturer’s protocol, 2x10^5^ to 2x106 D7 cells were subjected to CRISPR-Cas9/RNP nucleofection using electroporation enhancer (IDT, 10007805) and pairs of non-targeting control (IDT, 1072544 & 1072545), *IRF4*-targeting (IDT, Hs.Cas9.IRF4.1.AA, Hs.Cas9.IRF4.1.AB), or *PRDM1*-targeting (IDT, Hs.Cas9.PRDM1.1.AC, Hs.Cas9.PRDM1.1.AD) crRNAs pre-complexed with Cas9 tracrRNAs (IDT, 1072534). Cells were immediately recovered in complete RPMI-1640 media with an additional 10% FBS (20% final) and then added 1:1 to 2x stimulation media in 6-well plates for continued culture. Editing was confirmed by intracellular staining at 2, 7, or 14 days post-nucleofection, and indel efficiency at the *IRF4* and *PRDM1* loci was estimated by PCR & T7E1 endonuclease assay from genomic DNA harvested at D21 (14 days post-nucleofection).

### ELISpot assays

ELISpot assays were performed with 4h incubations at 37°C, 5% CO_2_ in clear-sided 96-well ELISpot plates (Millipore-Sigma, MAIPS4510) to enumerate the proportion of antibody-secreting cells (ASCs). Briefly, ELISpot plates were coated overnight (O/N) at 4°C with goat anti-Igκ & Igλ (1:1000 each; Southern Biotech, 2061-01 & 2071-01) diluted in 1x PBS. After washing, plates were blocked with complete RPMI-1640 media for at least 1h at RT, which was discarded immediately prior to addition of cells. 2:1 serial dilutions were performed in U-bottom microtiter plates and transferred to ELISpot plates, which were then incubated for 4hr at 37°C, 5% CO_2_. Plates were washed and incubated with goat anti-IgG (1:10000; Sigma-Aldrich, A3187-1ML) alkaline phosphatase (AP)-conjugated detection antibody diluted in 1x PBS O/N at 4°C in the dark. Plates were washed again and then developed for 3-10 minutes in the dark with BCIP/NBT-plus (MabTech, 3650-10).

### Flow cytometry and sorting

Flow cytometry gating for live surface phenotyping of human PBs (CD20^-^CD27^hi^CD38^hi^CD138^-^) and PCs (CD20^-^CD27^hi^CD38^hi^ICAM2^+^CD138^+^) was based on prior studies (Sanz et al., 2019; Staupe et al., 2022). Briefly, samples were washed with FACS buffer and incubated with Human TruStain FcX (BioLegend, 422302) for 20m at RT and then stained with antibody mastermix (50µL total) for 1h at 4°C in the dark. During the last 15min, Zombie Violet live/dead fixable viability dye (BioLegend, 423114) was added to the staining mix. Afterwards, samples were washed and resuspended in FACS buffer. For intracellular staining (ICS), samples were first live/dead stained, blocked, and washed as in surface staining, but then fix-permeabilized (eBioscience, 88-8824-00) for 20m at RT, blocked again, and then stained for 1h at 4°C in the dark. ICS samples were washed twice in 1x permeabilization buffer (eBioscience, 00-8333-56) then once in FACS buffer. ICS samples were further gated on non-debris to exclude background events after live gating. All analyses and sorts were performed on a 5-laser (R-Yg-B-V-UV) FACS Aria III using Single-Cell or Purity settings for activated B (CD20^+^CD27^-^) or PBs, respectively. Cells were sorted into complete RPMI-1640 media without phenol red, then diluted into FACS buffer for post-sort purity checks or washed twice for continued culture. For the sort-and-reculture experiment (Figure 1F, G and S1G), D7 CD20^+^CD27^-^ actB cells and D7 CD20^-^CD38^+^CD27^+^ PBs were sorted in parallel from the same culture and each returned to the standard D7-D14 then D14-D21 differentiation conditions described above, so that both populations were carried to D21 under identical conditions alongside an unsorted culture control.

| **Surface (live) flow cytometry mAbs (50µL final staining volume)** |  |  |  |  |  |
| --- | --- | --- | --- | --- | --- |
| Target | Fluorochrome | Dilution | Clone | Manufacturer | Cat. # |
| CD27 | PE-Cy7 | 1/50 | LG.3A10 | BioLegend | 124216 |
| CD38 | BV711 | 1/50 | HIT2 | BioLegend | 303528 |
| ICAM-2 (CD102) | FITC | 3/100 | CBR-IC2/2 | BioLegend | 328507 |
| CD138 | BV605 | 2/50 | MI15 | BioLegend | 356520 |
| CD19 | APC-eF780 | 1/50 | SJ25C1 | eBiosciences | 47-0198-42 |
| CD20 | BUV395 | 1/40 | 2H7 | BD Biosciences | 563782 |
| **Intracellular staining (ICS) flow cytometry mAbs (50µL final staining volume)** |  |  |  |  |  |
| Target | Fluorochrome | Dilution | Clone | Manufacturer | Cat. # |
| BLIMP1 | PE | 2/50 | 5E7 | BioLegend | 150006 |
| IRF4 | eF660 | 1/40 | 3E4 | eBiosciences | 50-9858-82 |
| IRF8 | PE-Cy7 | 1/40 | V3GYWCH | eBiosciences | 25-9852-82 |
| Ki-67 | BV786 | 3/100 | B56 | BD Biosciences | 563756 |
| CD138 | BV605 | 2/50 | MI15 | BioLegend | 356520 |
| CD19 | APC-ef780 | 1/50 | SJ25C1 | eBiosciences | 47-0198-42 |
| CD20 | BUV395 | 1/40 | 2H7 | BD Biosciences | 563782 |

### Flow cytometry analysis

Flow cytometry analyses were performed using FlowJo v10.9.0 software (Becton Dickinson and Company). Analysis gates were held constant between timepoints and donors for quantified populations (PB, PC). Biplot contours show 5% contour intervals, except for re-cultured cells in Figure 1F where biplots with 10% intervals are shown. Due to consistently low CD20^+^ frequencies compounding low cell numbers (<100 CD20^+^ live cells), D21 CD20^+^ cells from Donor 2 (Figure S1B, E, F) were excluded from visualization/analysis.

### Single-cell transcriptomics (scRNA-seq)

Cultured fresh (D19) or thawed (D7) cells were enriched for viable cells by Annexin V dead-cell depletion (STEMCELL Technologies, 17899) immediately prior to ß2m hash-tagging. Samples were hash-tagged using TotalSeq-C anti-ß2m (BioLegend, TotalSeq-C, see *feature_reference.csv* sheets in GEO submission for oligos) and prepared for 5’ Illumina kit GEX + hashtag oligo (HTO) sequencing as described by the 10x Genomics protocol *CG000149 (Rev D)* on D7 or D19 post-culture, or D9 (2 days post-nucleofection) for genomic perturbations, with a per-well target of 30,000 cells and 20,000 reads/cell. Reagents and hash-tagging antibodies were provided as sub-aliquots from the Single-Cell Sequencing Core (University of Pittsburgh). Sequenced libraries were aligned to the hg38 (GRCh38.p14; refdata-gex-GRCh38-2024-A) and HTO references using 10X Genomics Cell Ranger v8.0.0, then integrated and analyzed using the *Seurat* package (Hao et al., 2024) in R (R Core Team, 2021). Datasets were prepared for integration using the top 5000 variable features and corresponding principal components 1:50, then integrated using CCA-like `SCTransform(…, method=’glmGamPoi’, vst.flavor=’v2’, vars.to.regress=’percent.MT’, verbose=FALSE)`. 1-hot hashtag encoding was performed manually for each hash-tag using the following algorithm: 1) normalize all HTO counts using centered log-ratio (CLR) `NormalizeData(…,normalization.method=’CLR’, margin=2)`; 2) establish cutoffs emphasizing specificity over sensitivity based on violin plots; 3) assign HTO labels; 4) repeat steps 2 & 3 for each HTO until every cell is assigned hashtag(s); and 5) remove cells with 0 or more than 1 HTO. For D7/19 scRNA-seq analyses, empirical HTO cutoffs were: D7_1: CLR≥5.0, D7_2: CLR≥ 1.2, D19_1: CLR≥6.0, D19_2: CLR≥0.8, D19_3: CLR≥0.4. For D9 KO scRNA-seq analyses, empirical HTO cutoffs were CLR≥1.5 for all 6 HTOs (2 donors, 3 conditions). For analysis of the culture system to characterize prePCs, Leiden clustering was initially performed using `FindClusters(…,resolution=0.3, algorithm=4, method=’igraph’, n.start=30, n.iter=30, random.seed=15213).` For subsequent cluster marker gene identification, Model-based Analysis of Single-cell Transcriptomics (MAST) (Finak et al., 2015) was used with thresholds of log_2_(fold-change)≥0.40; minimum-fraction-expressing≥0.30; and adjusted p-value≤0.05. Low-quality (low-number/non-B-lineage/ambiguous) clusters were removed, with 11 remaining clusters. Each cluster was assigned as activated B cell (actB; *CD19*^hi^, *MS4A1*, *PAX5*, *IRF8*, *SPI1*, *SPIB*, and *EBF1*), plasma cell (PC; *PRDM1*^hi^, *IRF4*, *XBP1*, *CD27*, *CD38*, *SDC1*, *TNFRSF17*, and *JCHAIN*) or plasmablast/plasma cell precursor (PB/prePC; intermediate expression of actB- and PC-associated genes). After filtering, cluster annotation, and cluster aggregation, MAST was used to identify marker genes for each of the three major cell-types (actB, PB/prePC, and PC). The top 100 expressed DEGs for each cell type were used as gene modules using Seurat’s `AddModuleScore` function for scoring both BMPC-integrated and KO datasets. Cell-cycle scoring was performed using Seurat’s cell-cycle gene list (`cc.genes`) to infer G1, S, and G2/M phases. Within the PC cell state, isotype-specific DEGs were identified with MAST by comparing IgM vs. IgG/IgA, IgG vs. IgM/IgA, and IgA vs. IgM/IgG clusters as determined by *IGH* isotype heavy-chain expression. For KO scRNA-seq analysis, MAST (with the same thresholds as above) was used to identify *IRF4* and *PRDM1* KO DEGs based on the following comparisons: 1) *IRF4* KO PC DEGs: “Stunted PC” cluster versus the “PC” cluster, the former comprised nearly exclusively of *IRF4* KO cells, and 2) *PRDM1* KO PC DEGs: “*PRDM1* KO” PCs versus “Control” PCs, as non-actB *PRDM1* KO cells cluster with non-target control gRNA-treated PCs.

### TF ChIP-seq and GO enrichment

Enrichr (Kuleshov et al., 2016; Xie et al., 2021) was used for gene ontology (GO) enrichment of KO DEG sets (IRF4-only, BLIMP1-only, and co-regulated) in these 3 categories: ENCODE/ChIP-X Enrichment Analysis (ChEA), GO Biological Process, and GO Molecular Function. ChEA or GO terms were considered significant if adjusted p-value≤0.10 [-log_10_(adj. p-value)≥1].

### Single-cell transcriptional and chromatin accessibility profiling

Nuclei preparation for multiome-seq was performed as outlined in the 10x Genomics protocol *CG000365 (Rev C)*, with a per-well target of 10,000 nuclei with 20,000 GEX reads/nucleus and 40,000 ATAC reads/nucleus. 10X Genomics Cell Ranger Arc v2.0.2 was used to align GEX and ATAC data to the hg38 reference (refdata-cellranger-arc-GRCh38-2020-A-2.0.0). Paired GEX & ATAC data were processed, clustered, and integrated using ScanPy (normalizing counts using `log1p` and `1E04` total sum) followed by topic-modeling and joint low-dimensional embedding (MIRA) (Lynch et al., 2022). Pseudobulking based on integrated cell-states was performed for use in training ChromBPNet models used to identify putative IRF4- and BLIMP1-active genomic regions contributing to transcriptional activity (see below) (Pampari et al., 2025).

### ChromBPNet data analysis

Aggregated cell state-specific ATAC-sequencing reads were extracted from the single-cell multiome dataset by using Sinto’s `filterbarcodes` function. Resulting BAM files were used for peak calling with MACS2 `callpeak` function with following options: `--shift -75 --ext 150 --pvalue 0.01 --nomodel -B --SPMR --keep-dup all --call-summits -f BAMPE.` Peaks for each cell state were merged to generate a union set of ATAC peaks to train ChromBPNet models for each cell cluster. A Tn5 bias model was trained by running ChromBPNet `bias pipeline` with nonpeak sequences from inaccessible chromatin regions. Bias-corrected BigWig files were generated by running ChromBPNet `pred_bw` and base-pair resolution contribution score predictions were generated by running ChromBPNet `contribs_bw` with the union set of ATAC peaks. The resulting bigWig files were converted to bedGraph files using `bigWigToBedGraph` (ucsc_utilities). Profile contribution score tracks (i.e., predicted contribution to ATAC peak profile/shape) were used for all downstream analysis of ChromBPNet models.

### TF CUT&RUN

Whole cells were lightly fixed using 0.1% PFA for 10min, quenched with 1M molecular-grade glycine, and permeabilized using 1x eBiosciences Permeabilization Buffer (0.1% saponin final). Target-TF antibodies were tested for conformational binding by ICS flow cytometry. After fixation and permeabilization, steps detailed in the CUTANA ChIC/CUT&RUN Kit protocol were followed (Epicypher, 14-1048). Control (negative: IgG, positive: H3K4me3; included in kit) or target (IRF4: Invitrogen, 50-9858-82; BLIMP1: BD Biosciences, 564702) IgG antibodies were bound overnight at 4°C, and then 3-4x10^5^ cells were adsorbed to activated ConA beads. The assay was completed according to the CUTANA ChIC/CUT&RUN Kit protocol using 0.05% saponin instead of digitonin for subsequent buffers, with both 30min or 90min digestions for IRF4, BLIMP1, and IgG control, and 90min digestion for H3K4me3 control. Libraries were prepared using the CUTANA CUT&RUN Library Preparation Kit (Epicypher, 14-1001) and sequenced at the Health Sciences Sequencing Core (HSSC) at UPMC Children’s Hospital, targeting 12.5million reads per sample. 30min and 90min conditions were aligned using bowtie2 v2.4.5 (Langmead & Salzberg, 2012) with the following parameters: `--local --very- sensitive-local --no-mixed --no-discordant -I 20 -X 300 –dovetail`; then combined using samtools `merge` to achieve a target of 25 million reads per sample. Peaks were called from merged, deduplicated alignments using MACS2. Matched IgG and H3K4me3 normalized are included in the GEO submission as controls. Bedtools `intersect` was used to filter CUT&RUN peaks through OCRs from scMultiome, and to determine co-bound versus IRF4- or BLIMP1-bound OCRs for hypergeometric testing for enrichment.

### TF-bound OCR-to-DEG linkages

First, non-redundant ISREs (HOMER motifs 177, 179, 183) and EICEs (HOMER motifs 301, 302) were mapped to IRF4-& BLIMP1-bound OCRs from CUT&RUN peaks using HOMER’s `annotatePeaks.pl` (Heinz et al., 2010). Next, ISRE/EICE-containing OCRs were linked to genes if they were within 200kb of the TSS using GREAT (McLean et al., 2010). OCR-linked genes were checked for overlap with *IRF4*/*PRDM1* KO DEGs to identify putative direct targets (DEGs) of IRF4 and BLIMP1, then the OCR-linked DEGs were binned into: IRF4-only IRF4-bound DEGs, BLIMP1-only BLIMP1-bound DEGs, and co-bound co-regulated DEGs (concordant or discordant). DEGs that both TFs regulated at uniquely bound sites were excluded from systematic analysis in this study.

### Differential PWM analysis

Using the ISRE/EICE-motif hits, ISRE and EICE PWMs were regenerated for exclusively IRF4-bound or BLIMP1-bound sequences linked to uniquely regulated DEGs using seqinr (Delphine Charif, 2026), DiffLogo (Nettling et al., 2015), and Biostrings (Lifschitz et al., 2022). The Jensen-Shannon divergence was calculated and plotted as a logo using DiffLogo’s `diffLogoFromPwm` function using the regenerated PWMs.

### EMSAs

Cell-free IVT lysates were generated from reactions individually programmed with 3xFLAG-IRF4, HA-BLIMP1, 3xFLAG-IRF1, or HA-PU.1 expressed in a pT7CFE1-Chis vector using the 1-Step Human Coupled IVT Kit – DNA (Thermo Scientific, 88882). Lysates were stabilized with 5% glycerol and protease inhibitor and then stored at -80°C until ready for use. 32x 30bp, 5’ FAM-forward-strand labeled, HPLC-purified DNA probes (IDT) were synthesized or converted from the *CIITA* promoter context for EICEs and ISREs, respectively. Forward and reverse strands were resuspended in RNP Duplex Buffer (IDT, 11-01-03-01), heated to 90°C in a water bath for 5min, then slowly annealed 1:1 to 100µM final by allowing the water bath to cool to ambient temperature (∼2h). This stock was serially diluted to a working 20x concentration of 20nM, ultimately resulting in 1nM probe per binding reaction. 5x binding buffer was prepared using Tris (pH 7.4) with NaCl (50mM final), DTT, 5% glycerol final, and poly (dI:dC). 20µL binding reactions were incubated for 30min at RT. For supershift reactions, 1µL antibody was added 15min after beginning the reaction (anti-HA: Cell Signaling Technologies, clone C29F4 #3724S; anti-FLAG: Sigma-Aldrich, clone D6W5B #14793S). Gels were run at 100V, 1.5-2h, in the dark at 4°C and imaged with 10-minute exposures.

### Statistics

Flow cytometric or scRNA-seq comparisons of PB and PC frequencies across timepoints or perturbations were analyzed in GraphPad Prism v.10.5.0 (GraphPad Software, LLC) by two-way ANOVA with Tukey’s (Figure 1C, S1C), Sidak’s (Figure S2B), or Dunnett’s (Figure 3A, S3A) post-hoc tests, as indicated in the figure legends; n denotes independent experiments and error bars or ± ranges denote 1 S.D. For sort and re-culture experiments with actB and PBs (D7), PC-yield frequencies (D21) were log-transformed to achieve homoscedasticity (confirmed via F-test: p_raw_=0.0092, p_log_=0.4450) then analyzed using a one-tailed paired Student’s t-test (ratio paired t-test) to determine if D7 sorted PBs yielded more PCs than their actB counterparts. Differential gene expression was assessed with MAST using the thresholds given above, with Bonferroni-corrected p-values. Gene ontology and ChEA enrichments were considered significant at adjusted p-value≤0.10. Overlap between IRF4- and BLIMP1-bound OCRs was assessed by hypergeometric test for enrichment against all OCRs in R. Enrichment of discriminating N_1_/N_3_ motif variants at IRF4-versus BLIMP1-bound DEG-linked OCRs was assessed in GraphPad Prism by two-sided Fisher’s exact test, where categories were defined based on EMSA binding preferences (Figure 5A-D; counts in Table S8); two tests were performed and are interpreted with and without Bonferroni correction. Jensen-Shannon divergence between regenerated PWMs is reported qualitatively.

## Data Availability

Datasets generated by this study are deposited on NCBI Gene Expression Omnibus under GSE333613 (scRNA-seq), GSE333996 (scMultiome), and GSE333964 (CUT&RUN). Source data for the electrophoretic mobility shift assays are provided with this manuscript. Naive B cells were obtained from commercially sourced, de-identified healthy-donor PBMCs (STEMCELL Technologies); the study therefore did not constitute human-subjects research and read-level data were withheld to protect donor privacy.

## Acknowledgements

We thank Drs. Gina Doody and Reuben Tooze for their advice, including sharing a detailed human B cell culture protocol, and Dr. Garnett Kelsoe for the generous gift of MS40L^lo^ stromal cells used for B cell co-culture. We would also like to thank Drs. Shuxian Wu and Mark Shlomchik for their modified CUT&RUN protocol, which was adapted for the CUTANA CUT&RUN protocol. Furthermore, we appreciate the Unified Flow Core at the University of Pittsburgh for assistance with flow cytometry; the Single-cell Sequencing Core (Lafyatis Lab) at University of Pittsburgh for scRNA-seq sequencing and multiome library preparation; and Dr. William A. MacDonald from the Health Sciences Sequencing Core (HSSC) at Children’s Hospital of Pittsburgh for guidance with multiome and CUT&RUN sequencing. Last, we would like to thank Peter Gerges for logistical computational support, and Dr. Godhev Manakkat Vijay for experimental input and review of manuscript.

## Additional Information

This research was supported by grants from the UPMC ITTC fund and National Institutes of Health grants U01AI141990 (to H.S.), RO1AI145064 (to H.S.), and 5T32AI089443 (to N.A.P.). Additionally, this research was supported by a Cancer Research Institute Irvington Fellowship Award #4185 (to N.A.P.). Computational analyses were supported in part by the University of Pittsburgh Center for Research Computing and Data, RRID:SCR_022735, through the resources provided: the HTC and H2P clusters, which are supported by NIH award number S10OD028483 and NSF award number OAC-2117681, respectively. Schematic figures were made using BioRender.

---

**Figure S1.**
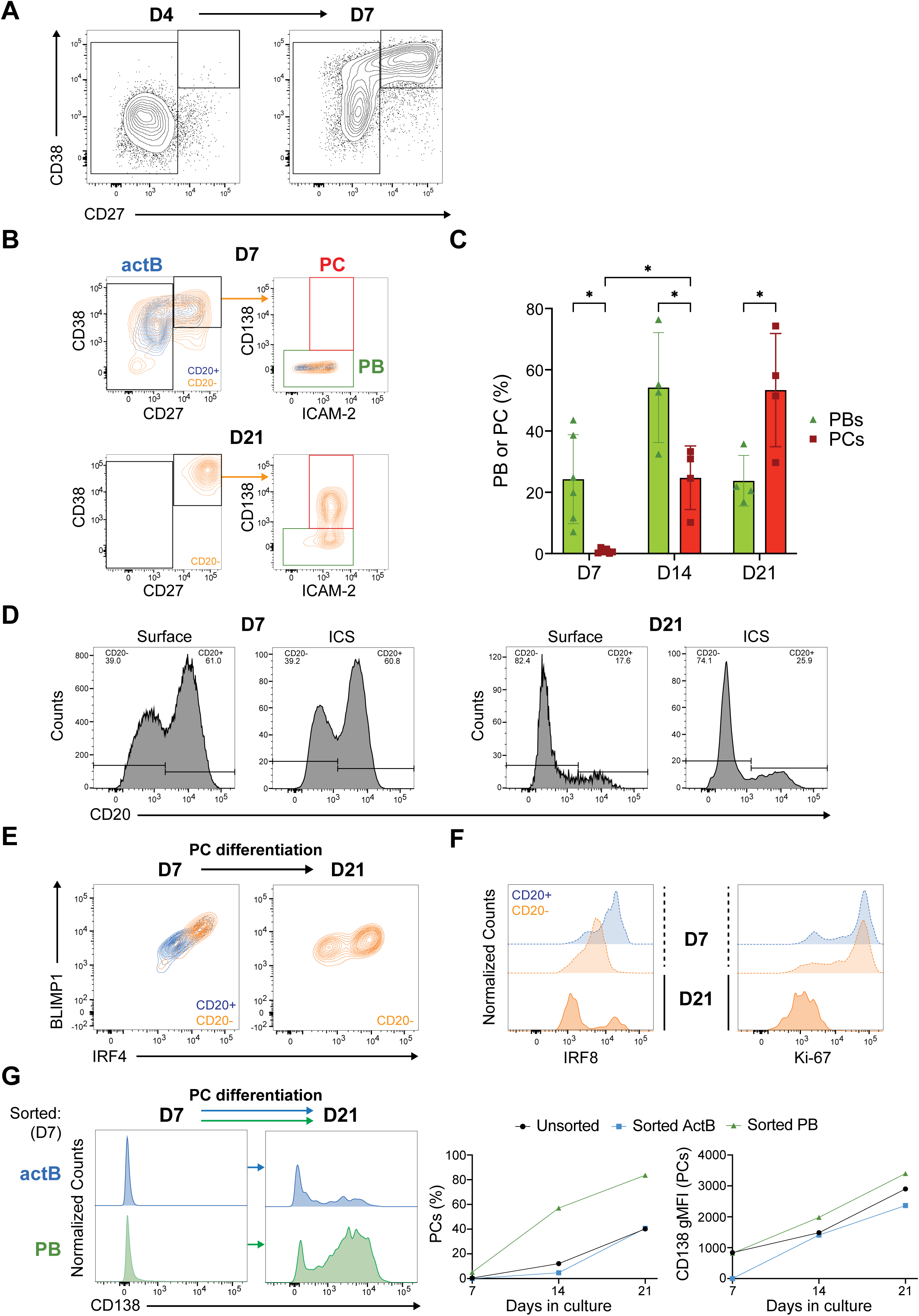
Stepwise human B cell differentiation generates PB/prePC and PC populations *in vitro*. (A) Upregulation of PB-associated CD38 and CD27 was monitored by flow cytometry at D4 & D7. Representative plots are shown for indicated timepoints. (B, C) PB and PC differentiation were monitored by surface immunophenotyping. (B) Representative flow cytometry plots show gating strategies used to quantify actBs (CD20^+^CD38^+^CD27^-^), PBs (CD20^-^ CD38^+^CD27^+^ICAM2^+/-^CD138^-^), and PCs (CD20^-^CD38^+^CD27^+^ICAM2^+^CD138^+^) at D7 and D21. (C) Bar graphs show PB and PC frequencies among live cells at the indicated timepoints (Donor 2). Dots indicate independent experiments, and error bars indicate mean +/- SD (n=4-6; *p<0.05, **p<0.01; two-way ANOVA with Tukey’s post-hoc test). (D) Histograms comparing CD20 staining and gating frequencies in surface-flow (Figure 1B and S1B) and intracellular-staining analyses (Figure 1D, E and S1E, F) at the indicated timepoints. (E, F) Intracellular flow cytometry was used to measure BLIMP1, IRF4, IRF8, and Ki-67. Cells were gated as CD20^+^ (blue) or CD20^-^ (orange) to relate intracellular measurements of the indicated TFs to the surface marker-defined PB/PC trajectory. (E) Biplots showing BLIMP1 and IRF4 expression at D7 (left) and D21 (right). (F) Histograms normalized to modal counts are shown for IRF8 and Ki-67 at D7 (dashed) and D21 (solid). (G) (Left) Histograms normalized to modal counts show surface CD138 expression of sorted D7 actB (blue) and PB (green) populations before and after their re-culture through D21. (Right) Quantification of D21 PC frequencies and CD138 gMFI within those PCs, stratified by condition: unsorted (black), sorted actB (blue), or sorted PB (green), related to Figure 1F, G.

**Figure S2.**
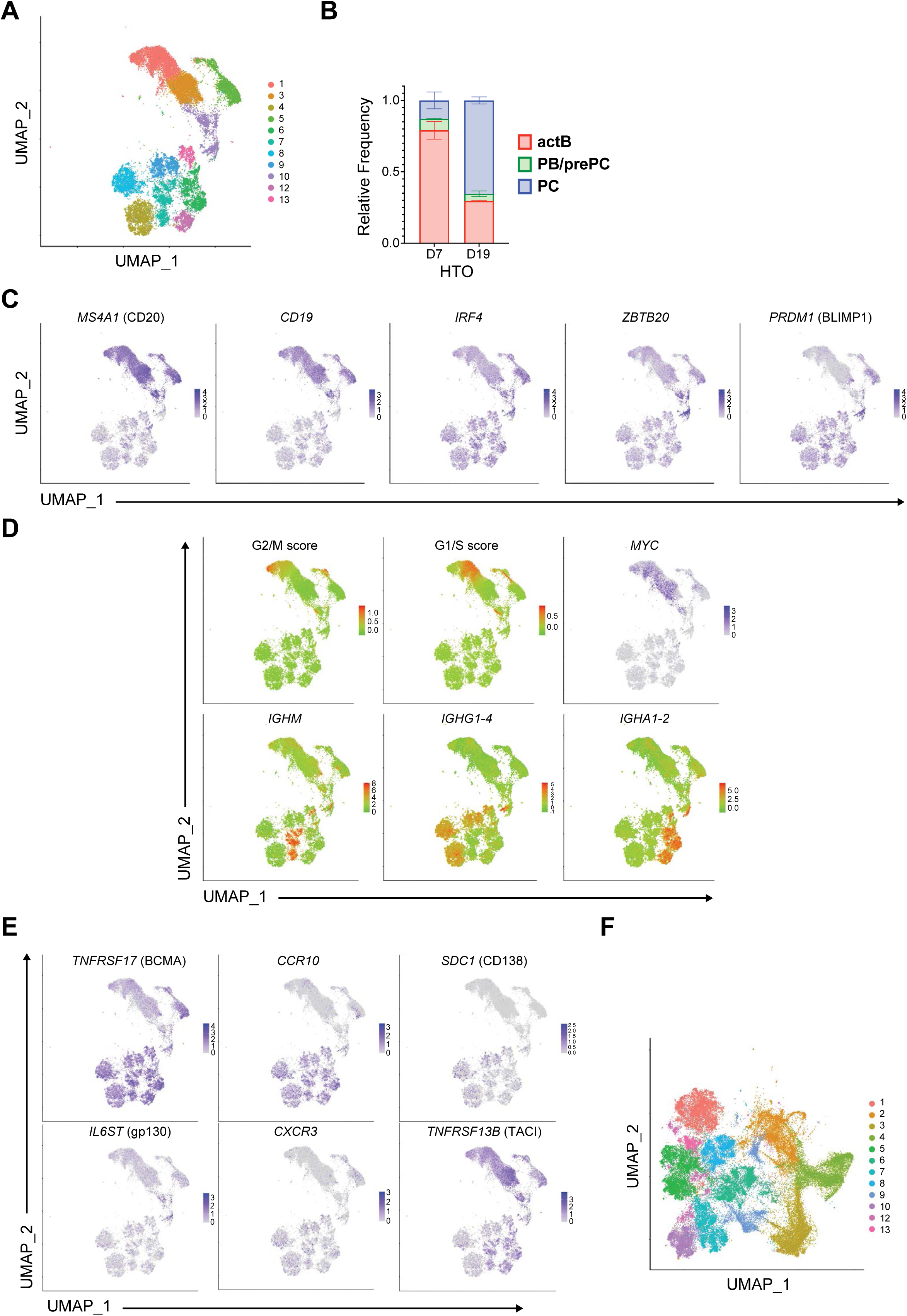
Single-cell transcriptional profiling resolves PB/prePC, PC, and isotype-associated states. (A) UMAP displays aggregated cells across timepoints (D7 and D19) and Donors 1 and 2, annotated by Leiden cluster. Donor 2 was assayed once at D19 to confirm mature PC transcriptome. (B) Bar plots show the relative frequency of each cell state at the indicated timepoints from Donor 1, where error bars denote S.D. (n=2 independent experiments; actB, p=0.0013; PC, p=0.0011; two-way ANOVA with Sidak’s multiple-comparison test). (C) UMAP projections show normalized expression of selected genes associated with PB/PC differentiation. (D) UMAP projections show cell-cycle module scores and *MYC* expression (top) and isotype scores derived from immunoglobulin heavy-chain transcript levels (bottom). Cell distributions are shown in Table S3. (E) UMAP projections show normalized expression of selected genes associated with PC homing and survival. (F) UMAP projection shows CCA-like integration of *in vitro*-derived cells and human BMPCs annotated by Leiden cluster.

**Figure S3.**
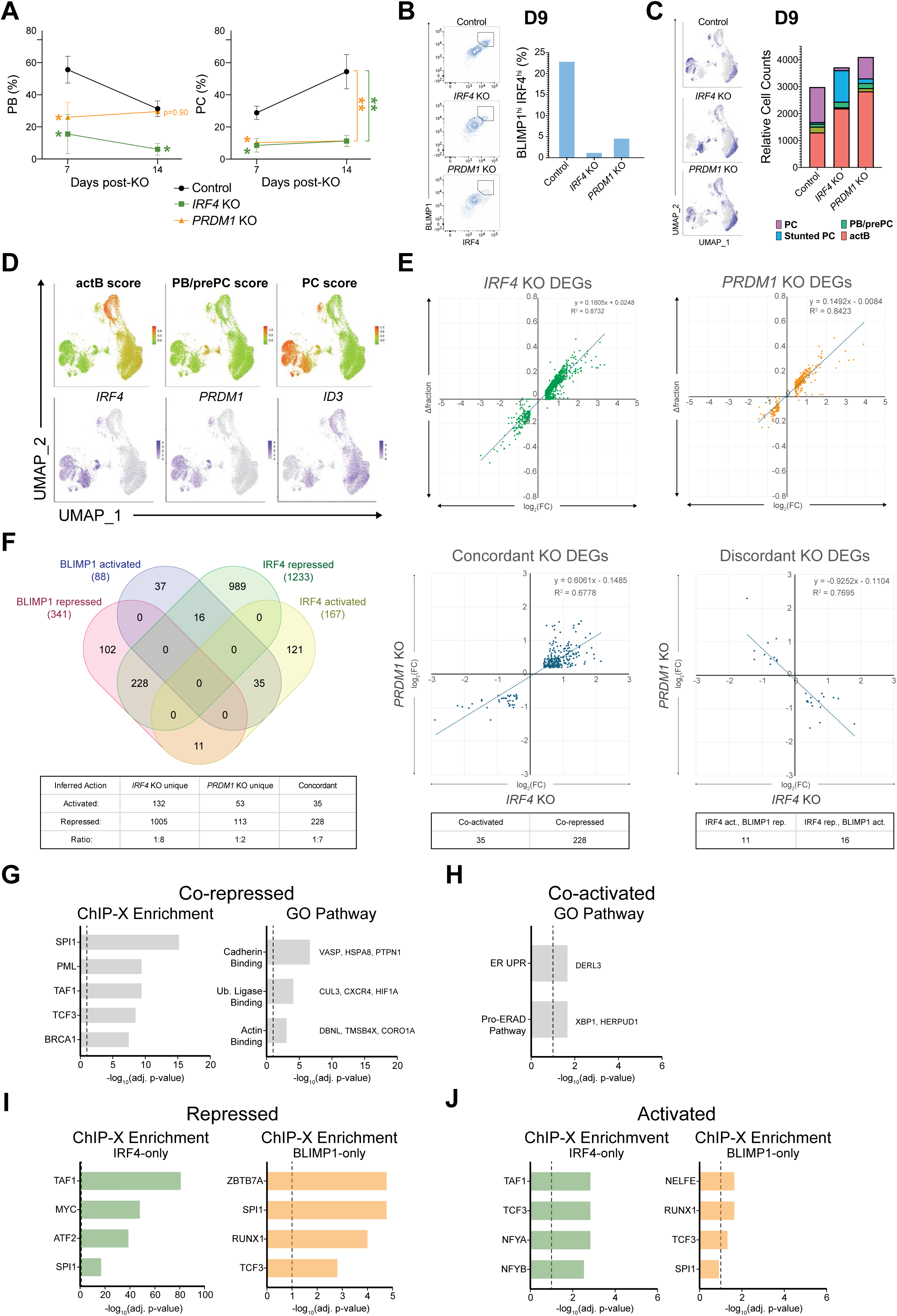
Stage-specific perturbations reveal distinct functions of IRF4 and BLIMP1 in prePCs. (A) Effects of stage-specific *IRF4* and *PRDM1* perturbations on PC differentiation were analyzed by immunophenotyping. Line graphs show mean PB and PC frequencies among live cells at D14 and D21 for Donor 2, with error bars showing S.D. (n=2 independent experiments; **p<0.01, ***p<0.001; two-way ANOVA with Dunnett’s post-hoc test for KO versus Control). (B) Effects of *IRF4* and *PRDM1* perturbation on IRF4 and BLIMP1 protein abundance were analyzed by intracellular flow cytometry. Representative plots show IRF4 and BLIMP1 levels at D9, 2 days post-KO (left). Bar plot shows BLIMP1^hi^IRF4^hi^ frequencies at D9 for Donor 2 (right), related to Figure 3B (representative of 2 experiments). (C) scRNA-seq analysis of Control, *IRF4* KO, and *PRDM1* KO cells from Donor 2 at D9. UMAP projections show aggregated cells annotated by Leiden cluster or sample identity from Donor 2, where gray cells are from Donor 1, related to Figure 3C. Bar plot shows the distribution of cluster cell numbers across Donor 2 samples at D9 (right). (D) UMAP projections show cell-state module scores derived from Figure 2A and Table S2 (top) and *IRF4*, *PRDM1*, and *ID3* expression (bottom). Cell-cycle score distributions can be found in Table S5, as was done in Table S3 for Figure S2D. (E) Correlation between log_2_(fold-change) and the change in fraction of cells expressing each DEG after *IRF4* or *PRDM1* KO relative to the indicated reference groups (*IRF4* KO: “Stunted PCs” versus “PCs”; “*PRDM1* KO” PCs versus “Control” PCs; see Methods). Amplitudes are shown as log_2_(fold-change) [KO – control], and frequencies are shown as the difference in fraction of expressing cells [KO – control]. (F) Venn diagram shows intersections among IRF4 and BLIMP1 DEGs, with unique and concordant DEG numbers summarized in the table below (left). Scatter plots show log_2_(fold-change) relationships for concordant (R^2^=0.68) and discordant (R^2^=0.77) KO DEG sets (right). (G, H) Bar plots show ChIP-X and/or gene ontology pathway enrichments for genes co-repressed (G) or co-activated (H) by IRF4 and BLIMP1, with representative genes indicated. (I, J) Bar plots show ChIP-X enrichment of target gene sets for IRF4-only or BLIMP1-only repressed (I) and activated genes (J).

**Figure S4.**
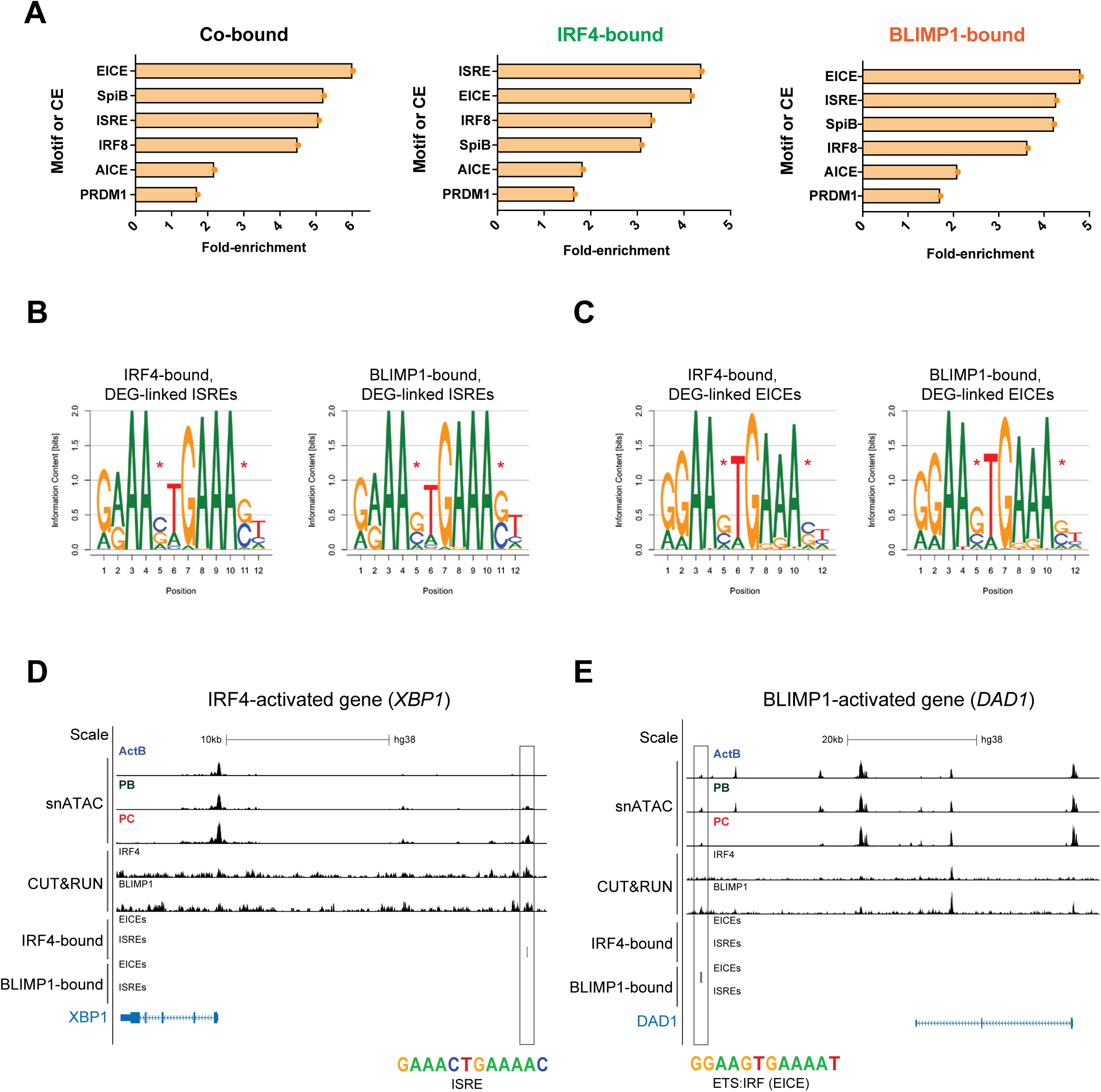
Differential IRF4 and BLIMP1 genomic occupancy reveals a discriminating ISRE/EICE motif lexicon. (A) Bar plots show motif enrichment analysis for selected IRF-related motifs in co-bound, IRF4-bound, and BLIMP1-bound OCRs. (B, C) Sequence logos show regenerated PWMs for (B) ISRE and (C) EICE motif instances at IRF4-bound OCRs linked to *IRF4* KO DEGs or BLIMP1-bound OCRs linked to *PRDM1* KO DEGs, related to Figure 4D and 4E. (D) UCSC Genome Browser tracks show an example of an IRF4-bound ISRE linked to the IRF4-activated gene *XBP1*. (E) UCSC Genome Browser tracks show an example of a BLIMP1-bound EICE linked to the BLIMP1-activated gene *DAD1*.

**Figure S5.**
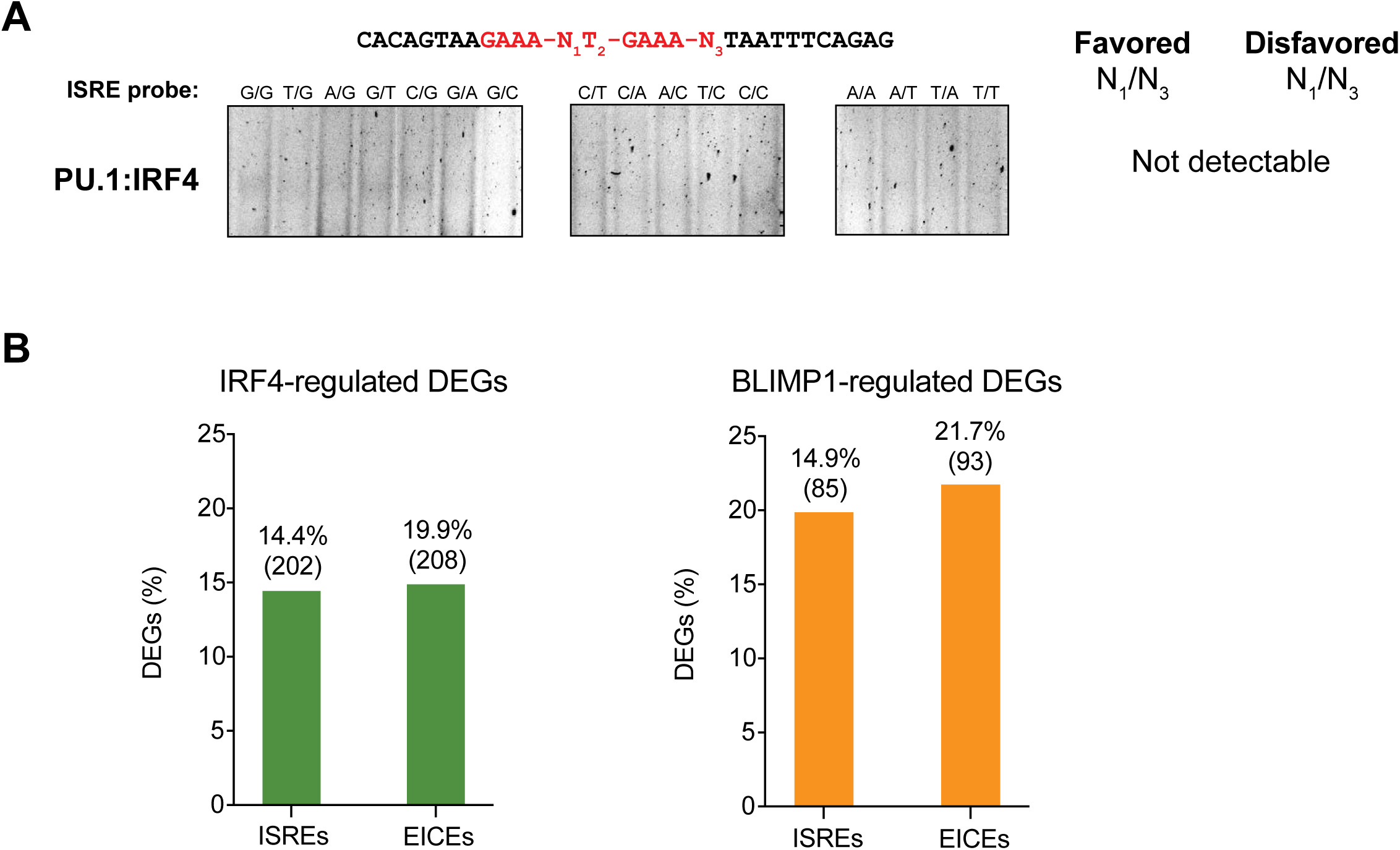
*In vitro* TF binding assays test base-pair rules and substantiate the ISRE/EICE regulatory logic. (A) EMSA analysis of IRF4 binding to ISRE probes in the presence of PU.1 instead of Ab directed against IRF4 epitope tag, related to Figure 5B. (B) Comparison of ISRE and EICE association with IRF4 and BLIMP1 DEG-linked OCRs. Bar plots show the frequency of IRF4 (left) or BLIMP1 (right) DEGs linked to IRF4-bound or BLIMP1-bound OCRs containing ISRE or EICE motifs. The number of DEGs in each category is shown in parentheses above the bar plots.

## Notes

### Competing Interest Statement

The authors have declared no competing interest.

### Summary of Updates

The manuscript has been revised in response to the public reviews at eLife.

