## Supplemental Table Legends for "Sequential and coordinated control of human plasma cell differentiation by IRF4 and BLIMP1 associated with a discriminating ISRE/EICE motif lexicon"

**Table S1. Flow cytometry population definitions and sort-and-reculture experiment statistics.** Definitions and gating hierarchy of surface-stained flow cytometry populations used for phenotyping or sorting. For the sort-and-reculture experiments, FACS-reported (D7) or manual hemocytometer (D14-D21) total cell counts are combined with flow cytometry frequencies to impute live cells and PBs, PCs, and non-PB/PCs within live cells. PC-generating yield per sorted population (A-D) is defined as the number of PCs generated at D21 per live D7 cell input. PC-generating yields are reported as mean ± S.D. of the two donors in the Results (one-way ratio Student's t-test $p=0.031$). Ratio of yields is defined as the mean PB/mean actB PC-generating yields (E, F).

**Table S2. Differentially expressed genes (DEGs) defining actB, PB/prePC, and PC cell states in scRNA-seq datasets.** DEGs were identified for the aggregated Leiden clusters expressing core B-cell or PC identity genes, with the resulting intermediate cluster labeled as PB/prePC (see Methods). DEGs are sorted in descending average $\log_2$(fold-change), with 2684 upregulated and 170 downregulated genes (total 2854 DEGs) defining actB; 814 upregulated and 834 downregulated genes (total 1648 DEGs) defining PB/prePC; and 138 upregulated and 2624 downregulated genes (total 2762 DEGs) defining PC. Of note, 20 of the 138 upregulated PC DEGs (14.5%) are Ig heavy-chain or kappa/lambda light-chain variable regions (*IGHV*, *IGKV*, *IGLV*). The actB, PB/prePC, and PC gene modules used for subsequent module scoring (Figure 2B and S3D) and heatmap visualization (Figure S2A) are comprised of the top 100 upregulated DEGs for each cell state. Each cell-state DEG set is separated as individual sheets/tabs.

**Table S3. Cell-cycle assignment and HTO cell-type distribution of *in vitro*-generated PCs and BMPCs.** Cell counts per Leiden cluster. Cell-cycle stage assignment based on Seurat's cell-cycle scoring function for each cell state (actB, PB/prePC, and PC). HTO distribution by cell state. Counts of isotype-specific PC clusters split by HTO based on dominant Ig heavy-chain (*IGH*) transcript.

**Table S4. DEGs of isotype-specific PC clusters.** DEGs were identified for the aggregated Leiden clusters expressing high *IGHM*, *IGHG1-4*, or *IGHA1-2* transcripts (see Methods). DEGs are sorted in descending average $\log_2$(fold-change), with 484 upregulated and 26 downregulated genes (total 510 DEGs) defining IgM PCs; 28 upregulated and 348 downregulated genes (total 376 DEGs) defining IgG PCs; and 100 upregulated and 24 downregulated genes (total 124 DEGs) defining IgA PCs. Each isotype-specific DEG set is separated as individual sheets/tabs.

**Table S5. Cell-cycle assignment and HTO cell-type distribution of *IRF4* and *PRDM1* KO scRNA-seq datasets.** Cell counts per Leiden cluster. Cell-cycle stage assignment based on Seurat's cell-cycle scoring function for each cell-type (actB, PB, PB/prePC, Stunted PC, and PC). HTO distribution by cell-type.

**Table S6. DEGs delineated by *IRF4* or *PRDM1* perturbations.** DEGs were identified for *IRF4* or *PRDM1* KO by comparing Stunted PCs versus PCs, or *PRDM1* KO PCs vs. Control PCs, respectively (see Methods). DEGs are sorted in descending average $\log_2$(fold-change), with 167 induced and 1233 repressed genes (total 1400 DEGs) defining *IRF4* KO effects; and 88 induced and 341 repressed genes (total 429 DEGs) defining *PRDM1* KO effects. Each KO DEG set is separated as individual sheets/tabs.

**Table S7. GO pathway and ChIP-X enrichment of inferred IRF4-only, BLIMP1-only, and co-regulated DEGs from indicated perturbations.** Enrichr was used to determine enrichment within ChIP-X Enrichment Analysis (ChEA), GO Biological Process, or GO Molecular Function categories (see Methods). Tables for each of 6 DEG sets are separated as individual sheets/tabs.

**Table S8. TF CUT&RUN summary and OCR-DEG contingency tables.** 1) Summary of CUT&RUN data: number and mean size of fragments (bp) with q-value cutoffs for MACS peak calling. 2) Summary of OCRs (multiome-seq) that are co-bound (overlapping), IRF4-only, or BLIMP1-only from CUT&RUN peak calls. Total OCRs from multiome-seq used to filter IRF4/BLIMP1 CUT&RUN peaks. 3) Contingency tables used for Fisher's exact tests to determine odds-ratio enrichment for IRF4-specific or BLIMP1 binding-competent EMSA-defined categories. 4) IRF4-bound or BLIMP1-bound OCRs (regardless of overlap) were scanned for strict ISRE/EICEs and associated with KO-specific DEGs within 200kb of TSS using GREAT, with numbers or % of bound ISRE/EICE-associated DEGs shown.
